# Maternal obesity and offspring neurodevelopment are associated with hypoxic gene expression in term human placenta

**DOI:** 10.1101/2024.07.10.602900

**Authors:** Fatima Gunter-Rahman, Shayna Mallett, Frédérique White, Pierre-Étienne Jacques, Ravikiran M. Raju, Marie-France Hivert, Eunjung Alice Lee

## Abstract

One third of women in the United States are affected by obesity during pregnancy. Maternal obesity (MO) is associated with an increased risk of neurodevelopmental and metabolic disorders in the offspring. The placenta, located at the maternal-fetal interface, is a key organ determining fetal development and likely contributes to programming of long-term offspring health. We profiled the term placental transcriptome in humans (pre-pregnancy BMI 35+ [MO condition] or 18.5-25 [lean condition]) using single-nucleus RNA-seq to compare expression profiles in MO versus lean conditions, and to reveal potential mechanisms underlying offspring disease risk. We recovered 62,864 nuclei of high quality from 10 samples each from the maternal-facing and fetal-facing sides of the placenta. On both sides in several cell types, MO was associated with upregulation of hypoxia response genes. On the maternal-facing side only, hypoxia gene expression was associated with offspring neurodevelopmental measures, in Gen3G, an independent pregnancy cohort with bulk placental tissue RNA-seq. We leveraged Gen3G to determine genes that correlated with impaired neurodevelopment and found these genes to be most highly expressed in extravillous trophoblasts (EVTs). EVTs further showed the strongest correlation between neurodevelopment impairment gene scores (NDIGSs) and the hypoxia gene score. We reanalyzed gene expression of cultured EVTs, and found increased NDIGSs associated with exposure to hypoxia. Among EVTs, accounting for the hypoxia gene score attenuated 44% of the association between BMI and NDIGSs. These data suggest that hypoxia in EVTs may be a key process in the neurodevelopmental programming of fetal exposure to MO.

## Introduction

The prevalence of obesity, both during pregnancy and generally in the population, is increasing^1^. 29% of women in America enter pregnancy with obesity, defined as a body mass index (BMI) >= 30 kg/m^2^ ^2^, and the prevalence of maternal obesity (MO) is expected to go up to 47% by 2030^3^. Similar trends are present in countries around the world^4^. MO is associated with life-long offspring mortality^5^ and morbidity, including cardiovascular disease, diabetes, cancer, psychiatric disorders^1^, and obesity^6^ . Studies show offspring DNA methylation patterns associated with MO at birth^7^, and later in childhood and adolescence^8^, suggesting that epigenetic programming plays a role in the lasting effects of MO.

Understanding the placental alterations related to MO can provide insights into the developmental programming of chronic disease, potentially serving as a basis for interventions. The placenta consumes 40-60% of oxygen and glucose delivered to the fetus, despite making up only 10-20% of uterine mass^9^, suggesting its importance. There is growing interest in the role of the placenta on neurodevelopment and mental health of the offspring later in life^10–13^, especially since placental gene expression and DNA methylation has been associated with offspring development of autism and schizophrenia^14,15^.

Until now, no cell-type specific profiling of the effects of MO on human placenta and trophoblasts has taken place. Previous studies have primarily examined changes in bulk gene expression^16–19^, and have not clarified a consistent placental face for sample site of origin. Gene expression within the same cell type differs by location within the placenta^20^. Single-cell RNA seq (scRNA-seq) or single-nucleus RNA-seq (snRNA-seq) can offer insights that are missed by bulk, and snRNA-seq in particular can offer insight into syncytiotrophoblasts (SCTs)^21^, since their multinucleated nature leads to poor recovery from scRNA-seq. To our knowledge, there is only one study that examined cell-type specific transcriptomic changes associated with maternal BMI in humans, but it was limited to maternal immune cell types in the decidua^22^, which misses some of the key cell types of interest in offspring development—including fetal macrophages (FM)^13,23^ and trophoblasts^24,25^.

Here, we elucidate cell-type specific transcriptomic changes associated with MO by generating snRNA-seq data from term placenta tissues of humans with a pre-pregnancy BMI of 35+ (MO condition) or 18.5- 25 (lean condition). We profile the maternal facing side and fetal facing side of the placenta separately. We also rely on offspring follow-up measures in the Genetics of Glucose regulation in Gestation and Growth (Gen3G) pre-birth cohort to understand how placental transcriptomic changes associated with MO relate to post-natal growth and neurodevelopment^26^.

## Results

### Single nuclei profiling of term placenta from MO and control pregnancies

We selected and received frozen term human placenta samples from the Women and Infants Health Specimen Consortium at the Washington School of Medicine in St. Louis, Missouri. We profiled nuclei from the maternal-facing side (3 lean and 7 MO samples) and fetal-facing side (4 lean and 6 MO samples) separately. All samples came from singleton pregnancies without pre-eclampsia or gestational diabetes (see Methods for full exclusion criteria). Samples in each condition and each side came from both fetal sexes. Maternal age ranged from 19 to 30, and did not differ significantly between the two groups on each side (maternal side students t-test p= 0.80, fetal side students t-test p=0.39). Additional demographic characteristics are in Table S1.

After filtering for low-quality nuclei (see Methods, Figure 1a) we recovered 62,864 nuclei. All major cell types were well-represented: trophoblasts, immune cells, and endothelial/ stromal cell populations, and sub-clustering was performed to identify detailed cell types (Figure 1a-b, e). Batches were well integrated (Figure 1c, S1), and placental cell types were represented on both sides, other than cell types known to reside in the decidua (Figure 1d, S2). We performed a sub-analysis on male fetuses to identify the origin (fetal or maternal) of cells found in the placenta. XIST is expressed only in females, and Y chromosome genes such as USP9Y are only expressed in males. Using these marker genes, we determined that immune cells, other than fetal macrophages (FM), were of maternal origin, trophoblasts were of fetal origin, and endothelial and stromal cells were mixed (Figure S3) as expected^20,27^.

**Figure 1:**
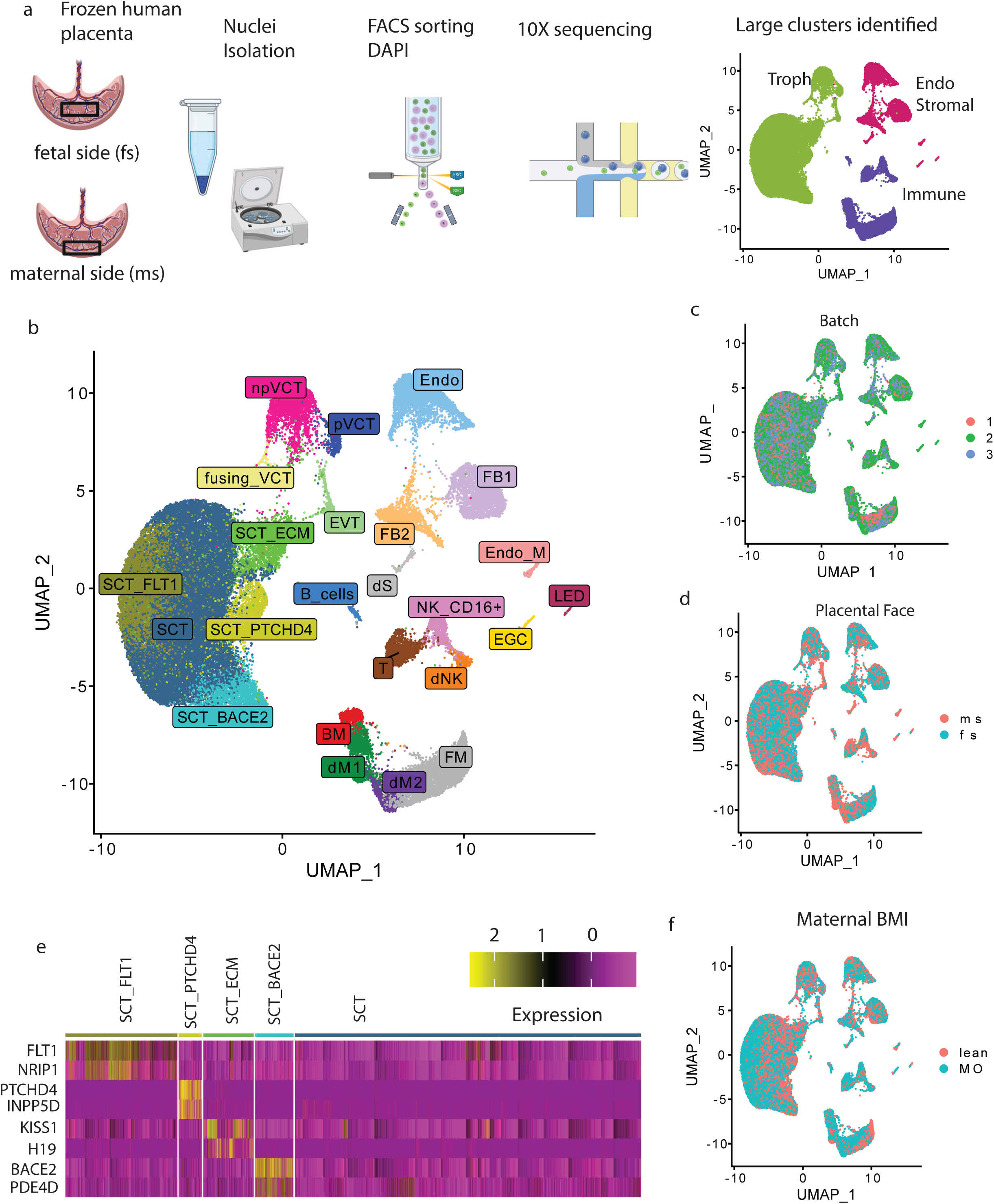
Overview of single-nuclei RNA-seq from human term placenta. 1a) Experimental procedures to generate single-nucleus data. b) UMAP plot of cell types identified. c) Batches are well integrated per UMAP colored by sequencing batch and d) most cell types have representation on both sides of the placenta. e) Marker gene expression of SCT subclusters identified. f) All cell types are present in both BMI conditions. Acronyms: SCT= Syncytiotrophoblast, ECM= Extracellular matrix, BACE2, FLT1, & PTCHD4 are marker genes; VCT= villous cytotrophoblast, (n)p=(non)-proliferative, EVT= extravillous trophoblasts; FB= fibroblast, Endo= endothelial, Endo_M= maternal origin endothelial cells; LED= lymphatic endothelial cell; EGC= epithelial glandular cell; NK= natural killer; “d” prefix= decidual; M= macrophage; FM= fetal macrophage; S= stromal cell

As our study used single-nucleus RNA-seq rather than single-cell RNA seq, we recovered more SCTs than previous human placenta studies^27–30^. Recently, Wang et al.^21^ identified several sub-clusters in SCTs based on early pregnancy and term multi-omics. Here, we corroborate some of their findings, by performing de-novo clustering and identifying sub-clusters enriched for expression of FLT1 (what they call “lSTB mature 2-a”), BACE2 (“lSTB mature 1-c”), and PTCHD4 (“lSTB nascent”). This suggests that these are true sub-clusters of SCTs present in human term placenta.

In addition, we identified another cluster whose marker genes were enriched for extra-cellular matrix remodeling (Figure S4), with genes such as KISS1 and H19 (Figure 1e). Although this cluster expressed genes that did not match any of markers identified in human placenta, snRNA-seq of mouse placenta identified an SCT sub-cluster (“SynTII”) enriched for cell–matrix interactions^31^. In addition, this cluster’s meaningful functional enrichment for a key process of SCTs suggests it may play a unique role. KISS1 is synthesized by SCTs, and interacts with the KISS1-receptor on extravilllous trophoblasts (EVTs) to influence their invasiveness^32^. Elevated KISS1 expression has been found in placenta of women with preeclampsia^33^. H19, a paternally imprinted gene^34^, is expressed by all trophoblast cell types, and also thought to play a role in EVTs angiogenic capacity^35^.

Each cell type and subtype identified was present in both MO and control pregnancies (Figure S5-6). We did not find a significant association between MO and cell type proportion (using scCODA^36^) .

### MO is associated with hypoxia in the placenta

We next performed analyses to identify differentially expressed genes (DEGs) by BMI in each cell type on each side separately. We compared MO samples to lean samples, accounting for biological and technical covariates (fetal sex, maternal age, delivery mode, and sequencing batch, see Methods). We relied on decoupleR^37^ to detect enrichment of DEGs in the 14 core pathways available in PROGENy^38^ assembled from a large compendium of publicly available perturbation experiments (Figure 2a-b). PROGENy outperforms other pathway enrichment methods and considers transcriptional targets of signaling cascades since post-translational modifications or other molecules involved in signaling might not show transcriptional changes^38^. Among all pathways, hypoxia was enriched among DEGs (FDR<0.05) in the most cell types on both sides. On the fetal side, hypoxia was most strongly enriched in DEGs of SCTs and SCT_FLT1 subtype (Figure 1a), and on the maternal side, hypoxia was most strongly enriched in DEGs of EVTs. Increased hypoxia was primarily associated with MO, although a few cell types on the maternal side showed enrichment in the opposite direction. Of the three cell types in which MO is associated with lower hypoxia, two (Endo_M and NK_CD16+) are of maternal origin (Figure S3). Together, these results show a strong hypoxia response of the placenta in the MO condition, primarily by cells of fetal origin.

**Figure 2:**
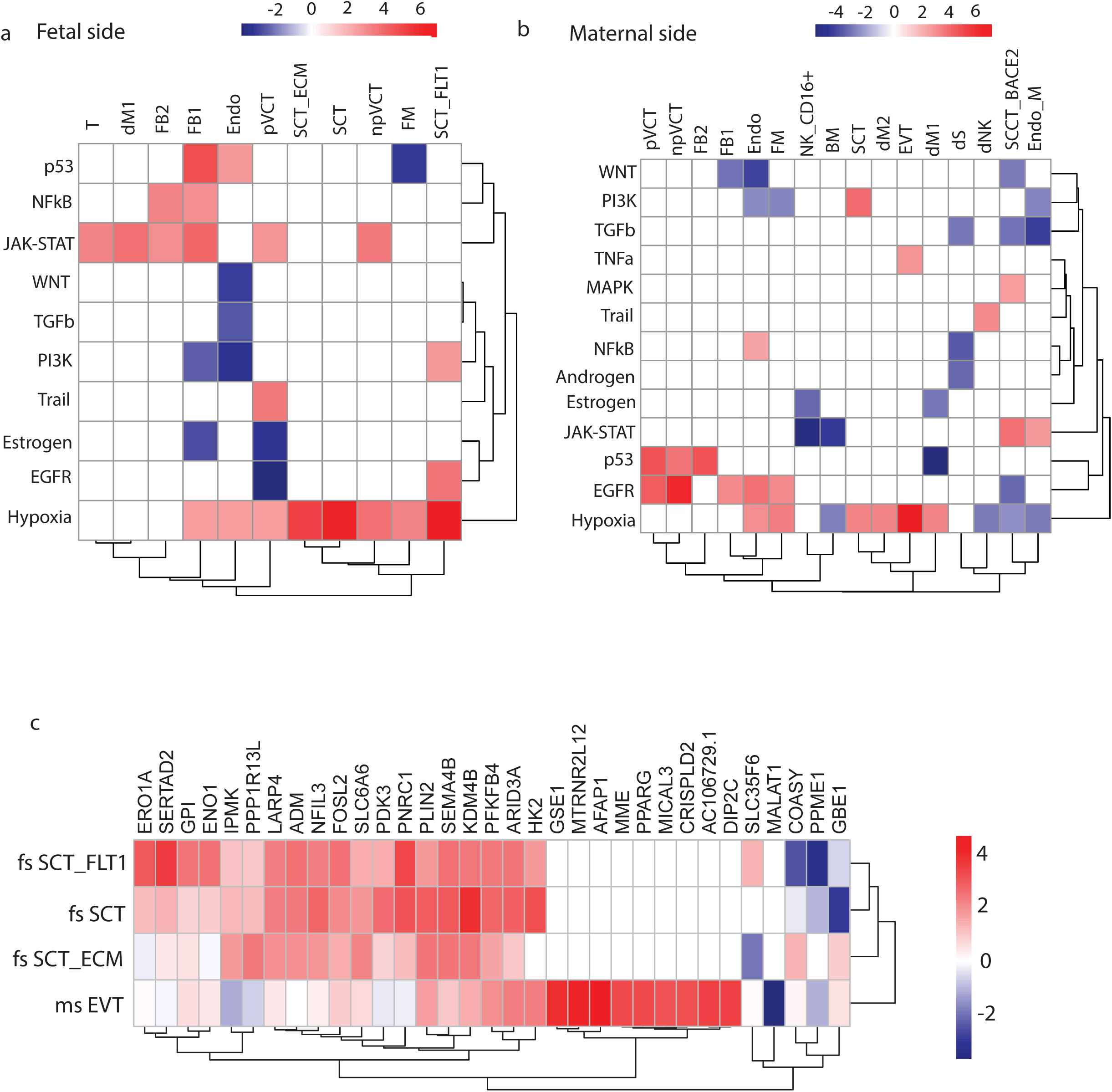
Maternal and fetal sides of placenta collected in pregnancies affected by MO are hypoxic. a-b) Pathway enrichment of genes differentially expressed by BMI on the fetal and maternal sides, based on PROGENy, showing enrichment for increased hypoxia response genes in multiple cell types. See Figure 1 legend for an explanation of cell types. All non-zero scores shown are significant with a false discovery rate (FDR) < 0.05. c) Genes from the hypoxia pathway in PROGENy and their differential expression by BMI, as represented by the DESeq2 test statistic, in four cell types (maternal side EVTs, fetal side [fs] SCTs, fs SCT_FLT1, fs SCT_ECM) with the highest enrichment for hypoxia. Genes were selected as the top 10 hypoxia genes associated with BMI in each of the four cell types. See Figure 1 legend for cell types and Table S3 for numerical results. For all (a-c) red represents positively associated with/ increased in MO, and blue represents negatively associated with/ decreased in MO.

We then explored which genes were relevant for hypoxia across the different cell types. We focused on the cell types whose DEGs by BMI had the strongest enrichment for hypoxia: maternal side (ms) EVTs, and fetal side (fs) syncytiotrophoblasts (SCTs) and two SCT subtypes on the fs: SCT_FLT1s, and SCT_ECMs (Table S2). For each cell type we identified the top 10 hypoxia associated genes, i.e. among the genes that are in PROGENy’s hypoxia pathway, the genes with the lowest p-value for association with maternal BMI for each cell type. This resulted in 33 genes across the four cell types, as there was some overlap.

We clustered the cell types and genes by their association with BMI (DESeq2 test statistic) and found that the ms EVTs were clustered separately from the rest. Although there were some genes with consistent direction of association with BMI across all cell types (Figure 2c), the top ten genes associated with hypoxia in ms EVTs were not expressed in the rest of the cell types (Table S3). These results suggest cell-type specific hypoxia responses, especially in EVTs.

### Hypoxia in the placenta is associated with short- and long-term offspring development

Because we observed hypoxia enriched among DEGs by BMI, we wanted to understand the biological relevance to the offspring of this hypoxia-related gene expression. We leveraged Gen3G pre-birth cohort with data from placental RNA-seq (from bulk tissue), in addition to birth outcomes, and child outcomes at ages 3 and 5 (Figure 3a, Table S4). We used PROGENy^38^ to calculate the hypoxia gene score (HGS) in samples from both sides of the placenta in Gen3G, based on a weighted sum of expression of hypoxia-associated genes (see Methods). We first investigated if the placental HGS was correlated with cord blood pH, as a reflection of fetal hypoxia^39^. If hypoxia is sustained, fetal production of hemoglobin can increase as a compensatory mechanism to increase oxygen delivery^40^, so we also investigated cord blood hemoglobin levels. We found that the HGS on both sides of the placenta was significantly correlated with cord blood PH and hemoglobin levels, and the correlations were stronger from the fetal-facing side biopsies (Figure 3b, Table S5). We also tested for a correlation between the HGS and Apgar score at five minutes post-delivery, as a measure of fetal distress and potential consequence of fetal hypoxia at the time of delivery. We again found a significant correlation with HGS on both sides, and a stronger correlation on the fetal side (Figure 3b, Table S5). Together, the correlations between the placental HGS and birth outcomes confirmed that it is a meaningful measure of biological hypoxia.

**Figure 3:**
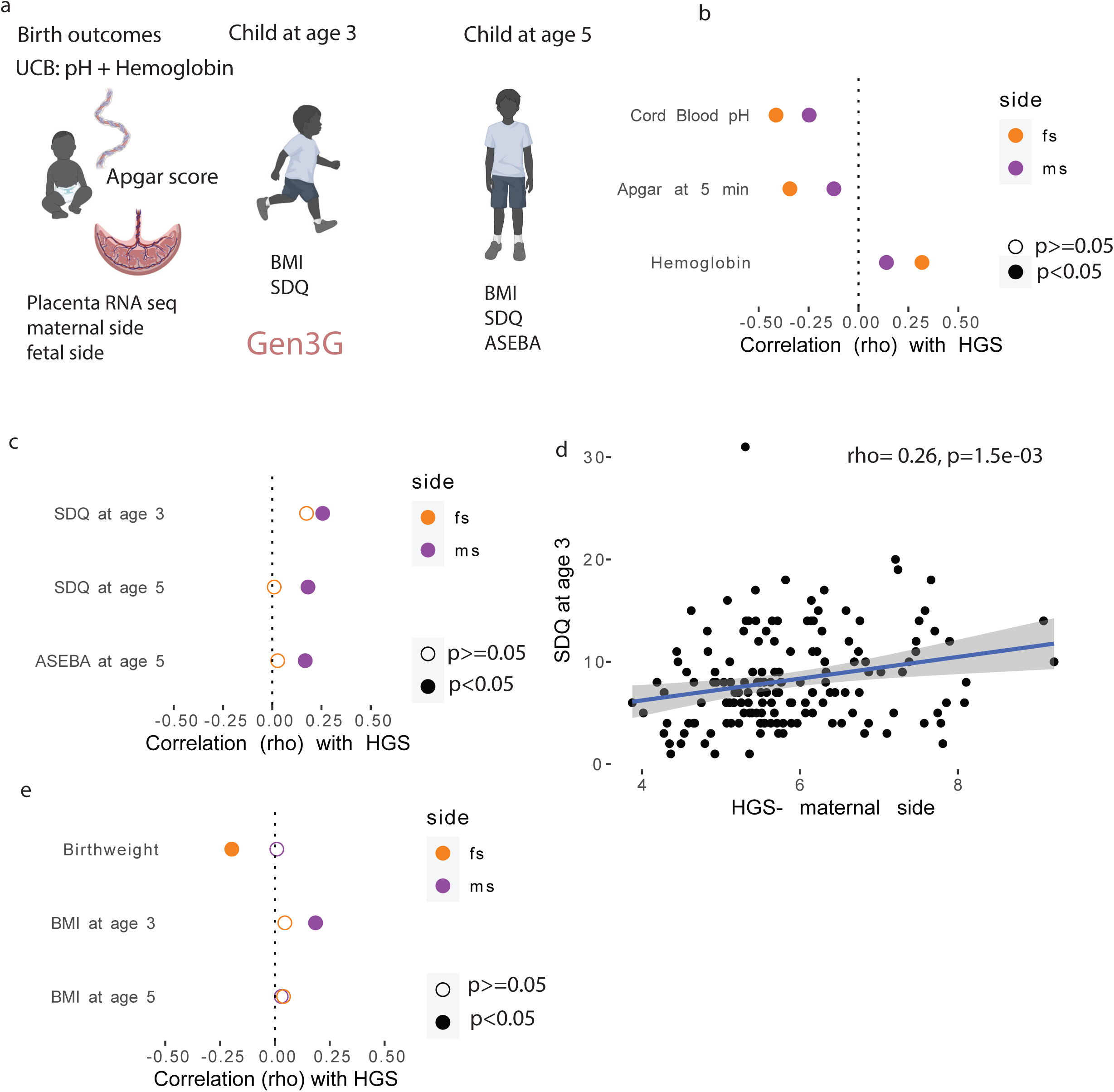
Hypoxia on the maternal and fetal sides have shared and unique correlations to offspring outcomes. a) Overview of characteristics profiled in Gen3G relevant to this study. UCB: Umbilical cord blood. b) Acute birth outcomes that are correlated with the placental hypoxia gene scores (HGS) derived from bulk tissue RNA-seq, with a stronger correlation on the fetal side. c) Placental HGS calculated from biopsies on the maternal facing side is associated with three neurodevelopmental outcomes: Strengths and difficulties questionnaire (SDQ) at ages 3 and 5 and Achenbach System of Empirically Based Assessment (ASEBA) at age 5. d) Visualization of the correlation between placental HGS from biopsies on the maternal facing side and offspring SDQ score at age 3. e) Correlations between placental HGS and growth-related traits.

We next examined childhood outcomes associated with higher maternal BMI in previous studies, namely impaired neurodevelopment^10,41^ and metabolic dysfunction^1^, for their relationship to the HGS. For neurodevelopment, we relied on two independent questionnaires: SDQ (Strength and Difficulties Questionnaire)^42^ and ASEBA (Achenbach System of Empirically Based Assessment)^43^. Higher scores on both the SDQ and ASEBA suggest more abnormal behaviors^42,43^. We found that higher placental HGS on the maternal side only was consistently correlated with higher scores of neurodevelopment (ND) testing at both ages 3 and 5, for all available measures (Figure 3c-d, Table S5). The fetal side of the placenta did not show a significant correlation for any of these ND outcomes. There was no consistent direction of association between childhood growth measures (birthweight, BMI at age 3, and BMI at age 5) of the offspring and hypoxia on either the maternal side or the fetal side (Figure 3e, Table S5), which led us to look further into ND-related gene expression.

### EVTs have the strongest signature of ND-related gene expression

To better understand the connection between hypoxia and ND, using Gen3G placental RNA-seq data, we performed a transcriptome-wide study to identify genes whose expression on the maternal side of the placenta was associated with eventual offspring SDQ score at age 3, correcting for key potential confounders such as maternal age and maternal education (see Methods, Figure 4a). We then similarly calculated genes whose expression was associated with offspring SDQ score at age 5, and ASEBA score at age 5. Several genes that were associated with the SDQ score at age 3 (FDR<0.05) were validated by their association with the SDQ score at age 5 and/or the ASEBA score at age 5 (FDR <0.05, same direction of fold change) (Figure 4b).

**Figure 4:**
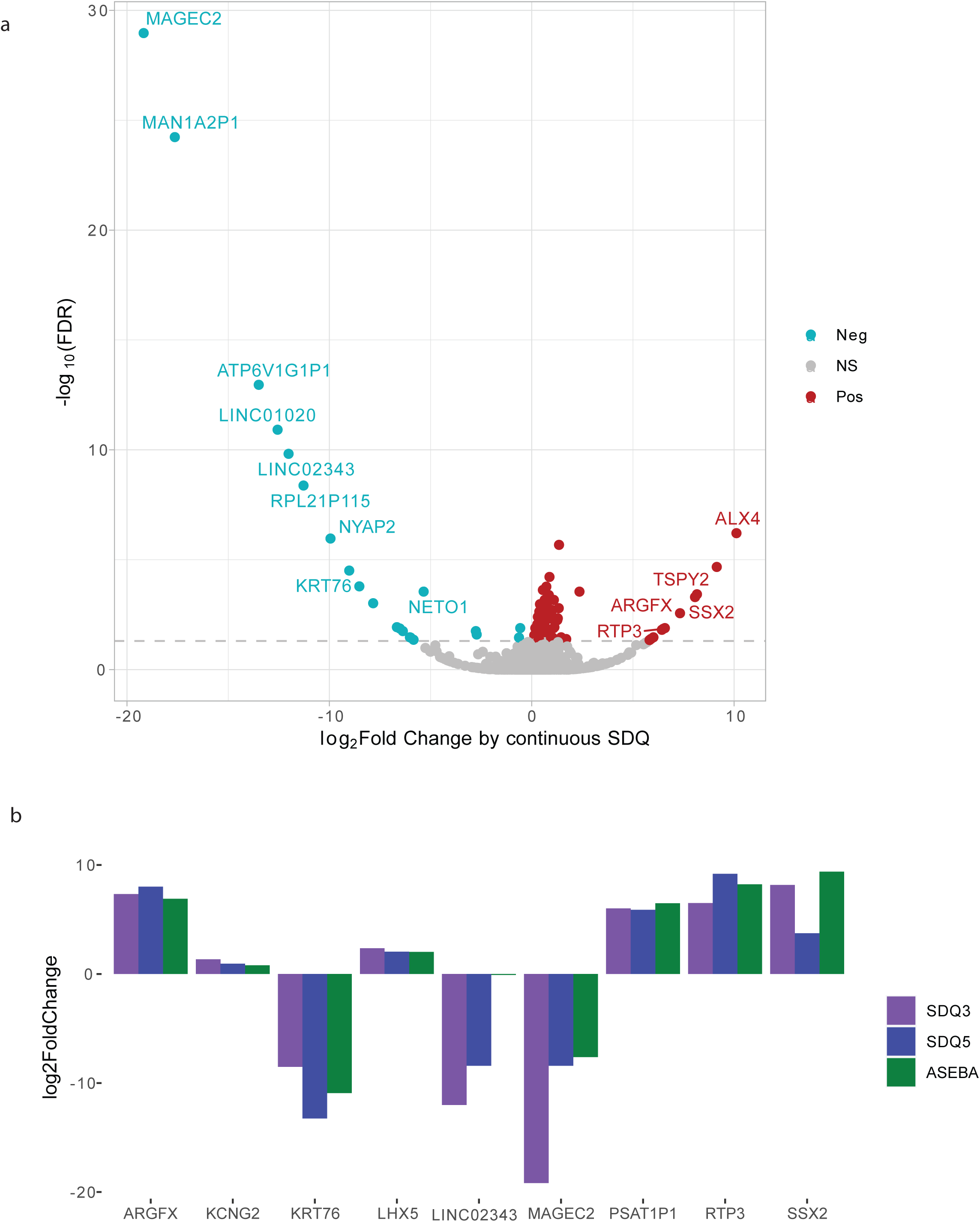
Genes whose expression on the maternal side of the placenta is associated with offspring neurodevelopment. a) Volcano plot of genes associated with SDQ score at age 3. Fold change represents the fold change in gene expression associated with a unit change in SDQ 3 score. b) Genes that were significant (FDR< 0.01) in a) and validated by also being associated with SDQ score at age 5 and/ or ASEBA score at age 5. SDQ= Strengths and difficulties questionnaire; ASEBA= Achenbach System of Empirically Based Assessment

We created placental neurodevelopmental impairment gene scores (NDIGSs), using the decoupleR^37^ package. The NDIGSs are a weighted sum of gene expression, with weights based on the association between placental gene expression and each ND outcome in Gen3G, such that genes positively associated with SDQ and/ or ASEBA scores in the offspring are positively weighted and genes negatively associated with the ND scores are negatively weighted. Genes with stronger associations with ND scores had larger (absolute value) weights (see Methods, Figure 5a). We calculated these placenta NDIGSs for all nuclei in the sn-RNA seq cohort, and grouped by cell type to identify cell types with the highest NDIGS. EVTs stood out clearly as the cell type with the highest NDIGSs for SDQ at age 3 (Figure 5b). The results ASEBA at age 5 were highly consistent (Figures S7). Although the results for SDQ at age 5 were noisier, EVTs still had high expression compared to other cell types (Figure S7). These patterns were present in nuclei from both MO and lean conditions (Figure S8).

**Figure 5:**
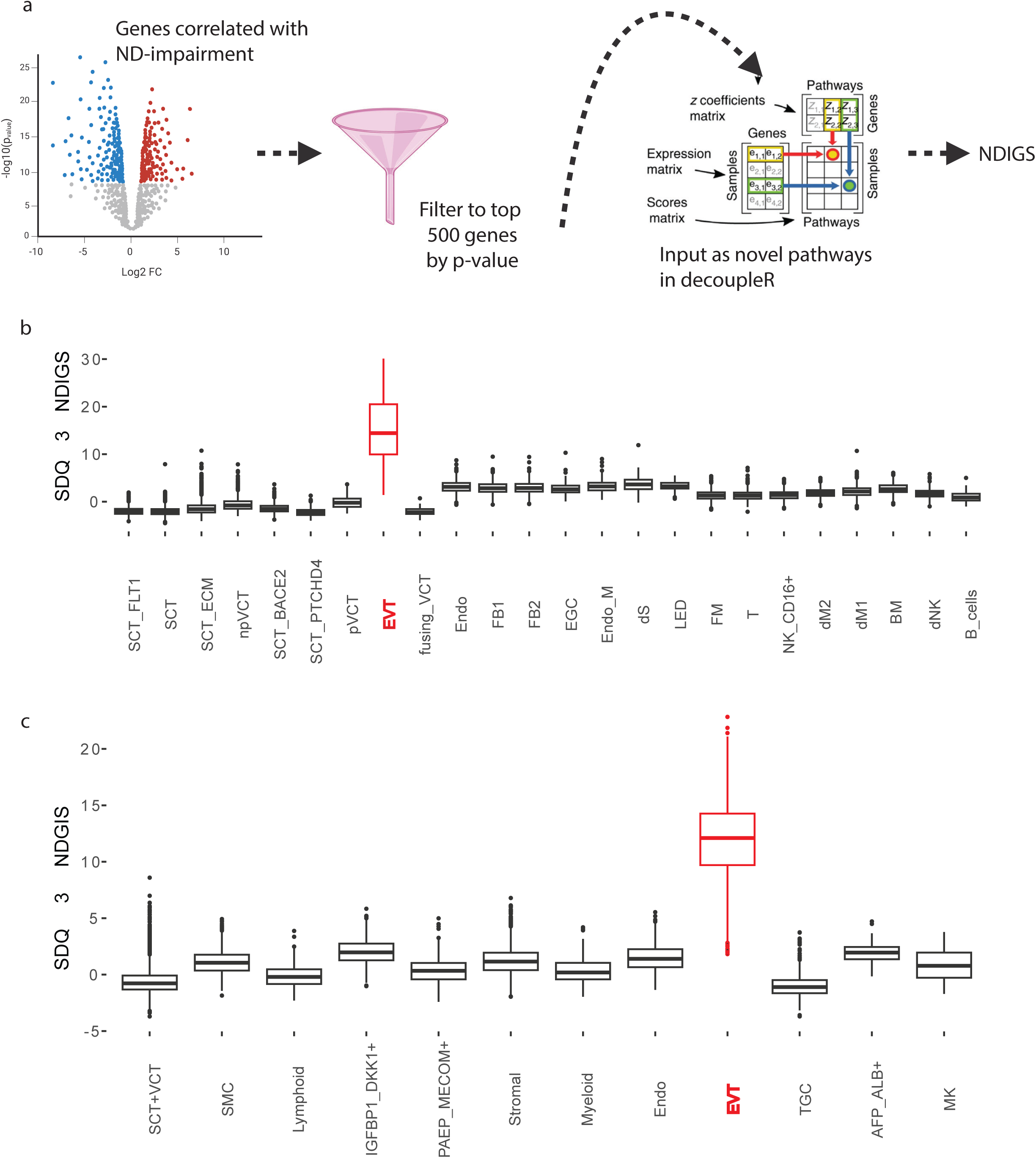
EVTs are a key cell type connecting hypoxia and neurodevelopmental impairment gene scores (NDIGS). a) Overview of generation of NDIGS scores for SDQ at age 3, SDQ at age 5, and ASEBA at age 5. b) SDQ 3 gene score by cell type in the single-nucleus dataset, showing the highest expression in EVTs. See Figure 1 legend for cell type acronyms. c) SDQ 3 gene score by cell type in a previously published second trimester placenta single cell dataset. The original cell types annotated by the authors are used. SCT, VCT, EVT, and Endo correspond to the same cell types identified in this paper. Lymphoid encompasses B cells, T cells, and NK cells. Myeloid encompasses all macrophages. MK= megakaryocytes. Please see Cao et al.^28^ for an explanation of the remaining cell types. SDQ= Strengths and difficulties questionnaire; ASEBA= Achenbach System of Empirically Based Assessment. For all box plots, the center line is the median, the upper box is the 75^th^ percentile, and the lower box is the 25^th^ percentile.

We turned to a single-cell dataset from second trimester placenta^28^ to see if EVTs had the highest expression of NDIGSs in an independent cohort, and if this finding was present at an earlier time point in gestation. EVTs again showed the highest expression across all cell types (Figure 5c), and this was consistent across all three NDIGS derived from SDQ at age 3, SDQ at age 5, and ASEBA at age 5 (Figure S9). This confirmed that EVTs have the highest expression of genes associated with offspring neurodevelopment.

### Hypoxia induces increased expression of NDIGSs in EVTs

We tested if placenta NDIGSs were correlated with placenta HGS in each cell type in both our term sn- RNA seq and the published sc-RNA seq from second trimester placenta^28^. EVTs stood out as having the highest correlation in our term single-nucleus dataset across all three NDIGSs (Figure 6a, Table S6). The correlation between HGS and NDIGS in EVTs was highly significant (Figure 6b; SDQ 3 rho=0.54 p<2.2e- 16, Table S6). This was also true for the second trimester dataset (Figure 6c, Table S7). Together, the results from two independent sc/sn-RNA seq datasets at different time points in gestation are consistent with the correlation found in Gen3G between HGS and ND outcomes, and goes further to identify EVTs as a key cell type in this correlation.

**Figure 6:**
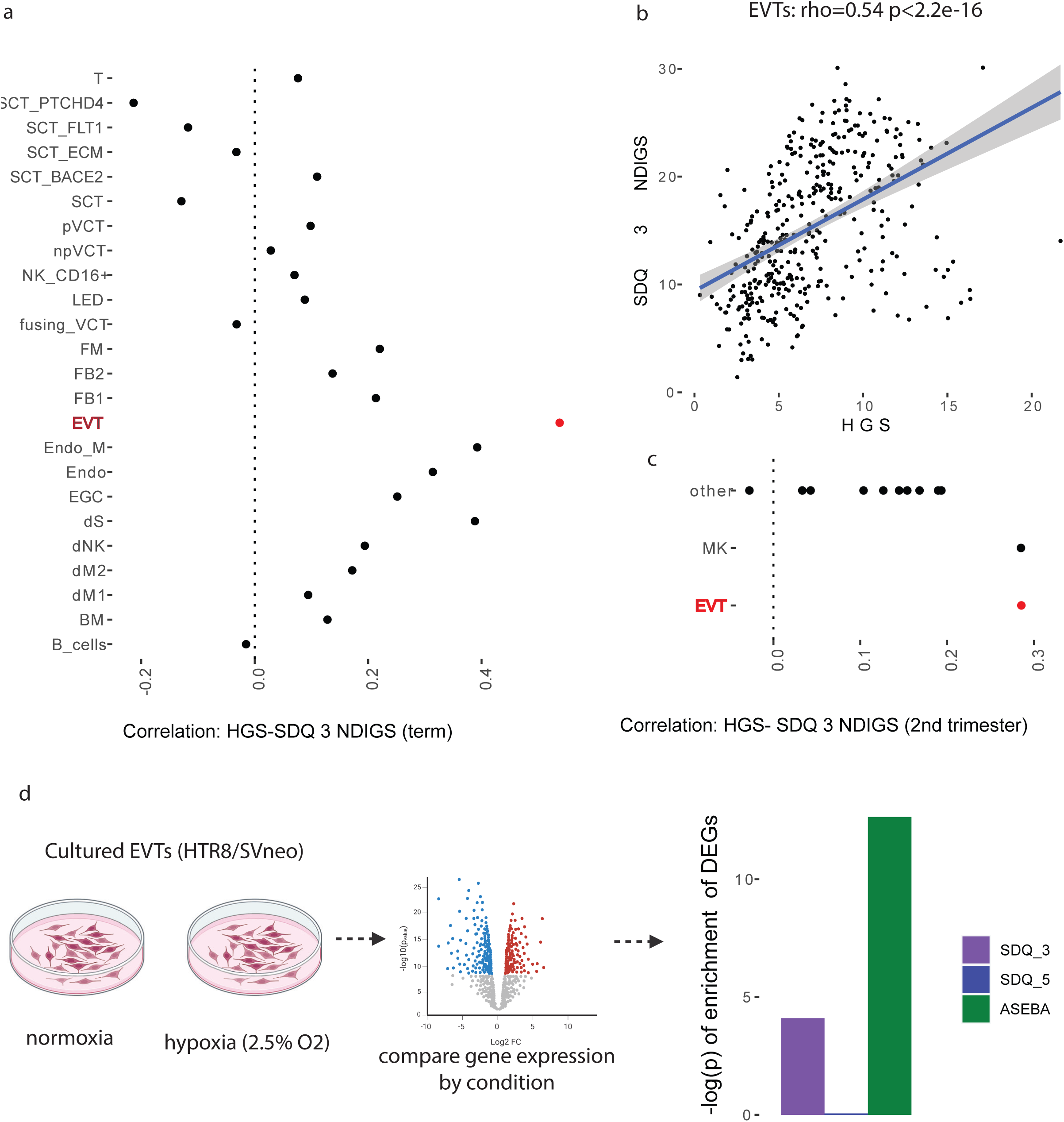
a) Correlations between SDQ 3 NDIGS and HGS across cell types in our term single nucleus RNA-seq. b) SDQ 3 gene score and hypoxia gene score are correlated in EVTs. c) Correlations between SDQ 3 gene score and HGS across cell types in previously published second trimester single cell RNA seq. MK= Megakarocytes. d) Exposing cultured HTR8/SVneo cells to hypoxia induces gene expression enriched for NDIGSs, reanalyzing previously published data^44^.

Given our findings in EVTs, we leveraged a dataset of HTR-8/SVneo cell line derived from invasive trophoblasts, an in vitro model of EVTs. Previous work exposed cultured trophoblasts to either hypoxia or normoxia conditions and performed bulk RNA-seq to calculate DEGs associated with oxygen levels^44^. We reanalyzed these DEGs and found a significant enrichment of genes included in NDIGSs (SDQ-3 enrichment: p= 7.9e- 5, ASEBA-5 enrichment: p= 2.3e-13, Figure 6d), with higher expression in the hypoxia condition (positive test score statistics of 3.95 for SDQ-3 and 7.34 for ASEBA-5). This shows that exposing human EVTs to hypoxia increases the expression of genes associated with impaired neurodevelopment, and supports our own observations.

### Hypoxia attenuates the relationship between BMI and ND score in EVTs

After exploring associations between placental gene expression scores of hypoxia and ND in EVTs, we return to maternal BMI, which initially highlighted hypoxia. Since EVTs were the cell type in which DEGs by BMI were the most enriched for hypoxia on the maternal side, it is possible that maternal BMI, placental hypoxia, and offspring ND are all connected. We first checked for enrichment of the NDIGSs among DEGs by BMI in the term sn-RNA data. EVTs stood out as the cell type whose DEGs by BMI had the most strongly significant enrichment for NDIGSs (Figure 7a). DEGs in fetal macrophages and endothelial cells also showed enrichment, albeit to a lesser extent. This shows that genes associated with MO are significantly enriched for genes associated with ND, and that this enrichment is strongest in EVTs.

**Figure 7:**
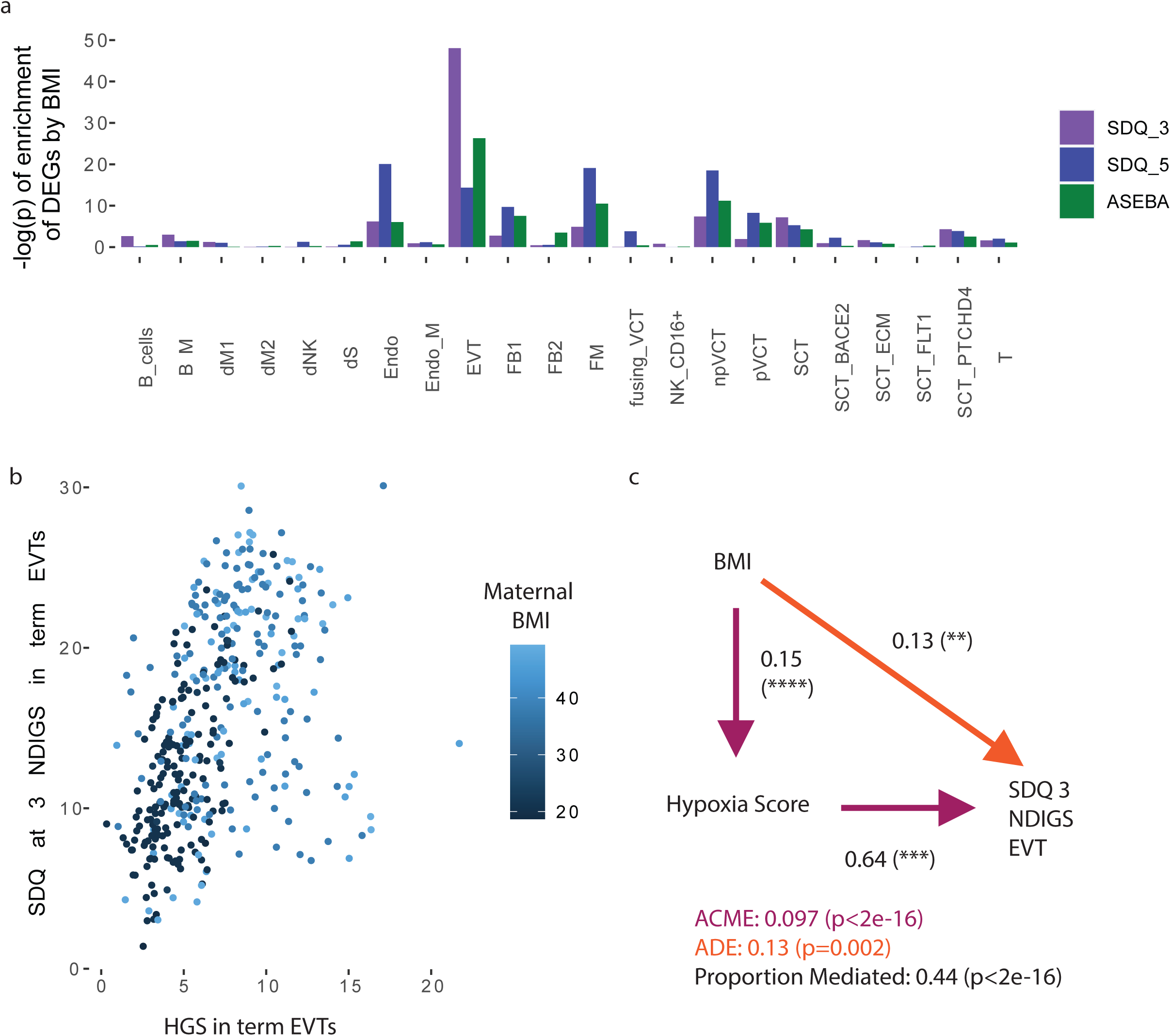
DEGs by BMI are enriched for genes associated with neurodevelopment, and hypoxia potentially links BMI to neurodevelopment. a) Enrichment of neurodevelopment-associated genes among each cell type DEGs by BMI, on the maternal side. b) Three-way relationship between BMI, hypoxia, and SDQ 3 gene score in EVTs. c) Accounting for hypoxia response gene expression attenuates the relationship between BMI and NDIGS scores in EVTs, for SDQ at age 3. ACME: Average causal mediated effect ADE: Average direct effect

We then examined the associations between maternal BMI, placental HGS, and NDIGS in EVTs, in the term sn-RNA data. Coloring the HGS-NDIGS plot from EVTs by maternal BMI shows that the nuclei with high HGS and NDIGS also come from mothers with a higher BMI than nuclei with low HGS and NDIGS (Figure 5b). We tested if accounting for HGS affected the relationship between BMI and ND score in EVTs, using the statistical mediation analysis^45^. We estimated the indirect effect of BMI on NDIGS through HGS, as well as the direct effect between BMI and NDIGS that remains when accounting for HGS, and found both significant indirect and direct effects (Figure 5c). 44% of the total association between BMI and NDIGS (SDQ at age 3) in EVTs was attenuated when accounting for HGS. The associations with SDQ and ASEAB at age 5 confirmed both a significant direct and indirect pathway linking BMI to NDIGSs, although the proportion attenuated by accounting for the HGS is lower (10% and 25% respectively). Together, this shows that a substantial proportion of the effect of BMI on neurodevelopmentally relevant genes in EVTs may be mediated by hypoxia.

## Discussion

In the current study, we showed that gene expression signature related to hypoxia in EVTs was associated with both maternal BMI and child neurodevelopment in humans. We rely on three independent cohorts to establish and validate our conclusions, including two single-cell and single-nucleus RNA seq datasets from different time points in gestation, and one large cohort with bulk RNA seq from hundreds of term placenta with offspring neurodevelopmental measures (Gen3G). The three cohorts are geographically and demographically diverse: Gen3G is a cohort of primarily White women in Sherbrooke, Quebec^26^; the single-nucleus RNA-seq generated in this study is from placenta of mostly Black women in St. Louis, Missouri, and the previously published second trimester dataset comes from the University of Washington^28^. The consistency of findings across these three cohorts increases the generalizability of our study.

Maternal BMI has been associated with fetal hypoxia as indicated by higher cord blood erythropoietin concentrations^46^ and hematocrit^40^, thought to be compensatory mechanisms. In mice, offspring of high-fat diet (HFD) dams have markers of oxidative stress in the placenta^47^ and, separately, in the fetal brain^48,49^, but the long-term consequences on the offspring of hypoxia associated with MO/ HFD have not been studied. Placental hypoxia, in the context of maternal uterine artery ligation in rodents, has been linked with structural changes in the offspring brain ^50,51^: reduced hippocampal size^52^, abnormal neural migration^53^, and delayed myelination^51,54^. We extend previous work by studying maternal BMI, placental hypoxia, and offspring neurodevelopment together, and by studying transcriptome-wide gene expression in term human placenta, rather than in animal models or cell culture.

Recent advances in single-cell technology have allowed for more easily profiling numerous cell types simultaneously. We generate one of the largest databases of transcriptomic profiling of human SCTs, and the only dataset to explore transcriptomic changes associated with maternal BMI in trophoblasts more generally. Our unbiased profiling of all cell types present demonstrated that EVTs consistently stood out as the cell type with the highest expression of NDIGSs and with the strongest correlation between HGS and NDIGSs. Given that EVTs are primarily on the maternal side of the placenta, this could explain the null relationship seen on the fetal side between HGS and neurodevelopmental outcomes in Gen3G. EVTs’ primary known function is to remodel the maternal spiral arteries in early pregnancy^55^, and their role in mid-to-late pregnancy is largely unknown. EVTs are highly sensitive to oxygen levels— hypoxia promotes EVT differentiation from stem cell precursors through HIF-dependent modulation^55^. It is possible that hypoxia, a normal condition in early pregnancy, aberrantly activates EVTs in the second and third trimesters, leading to changes relevant for neurodevelopment of the offspring. Synaptogenesis begins around week 16 of gestation continuing past birth, and myelination begins later, a few weeks before term delivery^41^. Key processes of neurodevelopment take place in the second half of pregnancy, and might be affected by oxidative stress.

Transcriptome wide association studies (TWAS) identified 139 placenta schizophrenia-specific risk genes, and of those predicted to be upregulated in schizophrenia, there was significant enrichment for expression in EVTs^14^. Emerging evidence from Mendelian randomization supports a causal role of trophoblast physiology (and inflammation from fetal macrophages) in the development of offspring depression^23,24^. In particular, EVTs and villous cytotrophoblasts were significantly enriched for genes whose variants are associated with offspring depression, and hypoxia is a postulated mechanism^24^.

Cultured trophoblast cells exposed to hypoxia release factors into media that caused damage to neurons, including decreased dendrite length. These effects on neurons are rescued by administration of MitoQ, an antioxidant^56^. In addition, fetal plasma from rats exposed to hypoxia during gestation contained miRNA that targeted genes expressed in the brain, and relevant for development^56^. There is also preliminary in vivo evidence that mitochondrial antioxidants can ameliorate the consequences of exposure to gestational hypoxia relating to offspring neurodevelopment^56^ and cardiovascular development^57,58^ in rodents. To add evidence for causality of hypoxia connecting maternal BMI to offspring outcomes, future studies should test if the use of antioxidants can partially rescue changes in the placenta and offspring brain associated with maternal BMI, using cell culture from lean and MO placental samples, or model organisms.

## Limitations of Study

Although the sc/sn-RNA-seq cohorts have tens of thousands of cells/ nuclei, they come from a small number of unique mother-child pairs. This limitation is in part compensated by corroboration of findings in the large bulk cohort, Gen3G. Since single-cell RNA seq is costly, future studies could perform bulk-RNA seq on sorted EVTs from more individuals to explore the roles of this cell type in a more cost-effective way. As a primarily observational study with human tissue, we are unable to prove causality, beyond the increase in neurodevelopmental gene expression following cultured trophoblast’s exposure to hypoxia.

## Methods

### Generation of single-nucleus RNA seq data

Human placenta samples were obtained at Washington University in St. Louis, as a part of the Women and Infant Health Specimens Consortium, following institutional review board approval. For this study, we selected term, singleton pregnancies that were not complicated by any maternal autoimmune disorder, active infection, substance abuse, preeclampsia, gestational hypertension, or placental abruption. Additionally, no mothers had pre-existing or gestational diabetes, and all mothers were less than 35 years old. All samples in the control category had a pre-pregnancy BMI within 18.5-25 and all MO samples had a BMI greater than 35. Samples were stored frozen at -80° C until use. The maternal side was sampled by slicing a thin 10mm layer off maternal facing side of the placenta (including basal plate), following peeling back of the chorioamniotic membranes. The fetal side was sampled by taking the spongy tissue located directly under the basal plate, making sure to exclude any basal plate.

Nuclei were isolated from placental samples following a modified version of the Allen Brain institute protocol for nuclei extraction. Briefly, tissue was homogenized using an electric pulsator and via dounces in a glass tissue homogenizer. A series of two washes was used to remove debris, and then nuclei were stained with DAPI. Flow cytometry was used to count 10,000 nuclei per sample, with gating based on DAPI-positive cells. The standard Chromium 10X v3.1 kit instructions were followed to generate the single-nucleus RNA-seq data. CellRanger was used to align sequencing reads to hg38 (CellRanger 6.0.1 for the four pilot samples in batch 1, and 6.1.2 for the rest of the samples. All batches had representation from both MO and control groups). CellBender 0.3.0^59^ was used to remove ambient noise from droplets with nuclei, and to identify empty droplets. Seurat 4.4.0 ^60^ was used to perform filtering (gene count >200, UMI count> 500, mitochondrial reads percentage <5), integrate samples accounting for batch, and perform clustering. scDblFinder 1.15.4^61^ was used to identify doublets, and all nuclei called as doublets were removed, as well as clusters that were dominated by doublets. Initial clustering identified 3 major cell types (trophoblasts, immune cells, and endothelial-stromal cells).

Within each large cluster, nuclei were re-integrated and re-clustered to improve performance and identification of rarer cell types. singleR 2.2.0^62^ was used to map to existing clusters from Vento-Tormo et al.^27^, and Seurat FindMarkers was used to identify markers of novel SCT subclusters. Markers were compared with Wang et al.^21^. gProfiler^63^ was used to calculate pathway enrichment of SCT sub-clusters.

### Pathway enrichment among DEGs in sn-RNA seq

Pseudobulk counts were calculated for each cell type on each side of the placenta by summing counts for all relevant nuclei of an individual. For each cell type and placental side, genes with at least 10 counts across all samples were kept, and genes with less than 10 counts were removed for low expression.

Differentially expressed genes (DEGs) by maternal BMI were calculated using DESeq2 1.40.2^64^, including maternal age, fetal sex, delivery mode, and batch as covariates for each cell type on each placental face. decoupleR 2.6.0^37^ was used to calculate enrichment of the 14 canonical pathways from PROGENy^38^. We followed the guideline’s in the tutorial (https://saezlab.github.io/decoupleR/articles/pw_bk.html). We relied on the multivariate linear model (mlm) tool on the DESeq test statistic, and used the top 500 genes for each pathway in PROGENy. False discovery rate^65^ (FDR) corrections were applied to correct for multiple hypothesis testing. pheatmap^66^ was used to make all plots.

To calculate enrichment of neurodevelopment related genes among BMI DEGs, we calculated novel gene weights (see Calculating genes whose expression in placenta is associated with neurodevelopmental impairment), just like those that exist in PROGENy^38^ for pathways such as hypoxia. The univariate linear modeling (ulm) tool from decoupleR was used to test for enrichment of these neurodevelopmental impairment genes scores (NDIGSs) among the BMI DEGs, relying on the test statistic from DESeq2.

### Gen3G cohort

Gen3G is a pre-birth cohort of pregnant women at the Centre Hospitalier Universitaire de Sherbrooke (CHUS), Quebec (Canada). Participants entered the study between 1 January 2010 to 30 June 2013.

Multiple pregnancies and women with regular use of medications that influence glucose regulation were excluded from the cohort. All study participants provided informed written consent, and the study protocols were reviewed by the ethical committees from CHUS, and from the Harvard Pilgrim Health Care Institute. To most closely match the single-nucleus RNA seq cohort, in this study, we excluded Gen3G mother-child pairs with maternal smoking during pregnancy, preeclampsia or gestational diabetes, birth before gestational age of 37 weeks, babies born small for gestational age, or placentas that were manually evacuated.

Birthweight was measured using standard clinical procedures within 2 hours of delivery. Cord blood and one cm^3^ of placental tissue from maternal and fetal facing sides were collected at delivery. For more information on the Gen3G cohort and biospecimens collected at birth, please see Guillemette et al.^26^ Total RNA was extracted from placenta, collected at the time of delivery. Samples with an RNA integrity of at least 4 were sequenced (n=466). For more details on pre-processing of Gen3G RNA-seq data, please see the “RNA extraction, sequencing and QC” section of Hivert et al.^67^

BMI was measured in the children at ages 3 and 5. Mothers completed the Strengths and Difficulties Questionnaire (SDQ)^42^ for their children at ages 3 and 5, along with the Child Behavior Checklist from the Achenbach System of Empirically Based Assessment (ASEBA)^43^ at age 5. For more details on the neurodevelopmental measures in Gen3G, please see Faleschini et al.^68^

### Hypoxia gene score correlations in Gen3G

The hypoxia gene score (HGS) was calculated by applying progeny to the expression matrix of all placental samples in Gen3G that met exclusion criteria described above. mlm from decoupleR was used to calculate an enrichment score of each pathway in PROGENy (including hypoxia). The Shapiro-Wilk normality test (shapiro.test, base R 4.3) was used to test for normality of hypoxia scores among placental samples on the maternal side and fetal side.

Hypoxia scores on both sides were not normally distributed per the Shapiro-Wilk normality test^69^ (fetal side p=6.8e-04, maternal side p=5.2e-07), so we used Spearman correlations to test for an association between HGS and cord blood pH, fetal hemoglobin, Apgar score at five minutes, birthweight, neurodevelopmental scores at ages 3 and 5, and offspring BMI at ages 3 and 5.

### Calculating genes whose expression in placenta is associated with neurodevelopmental impairment

We used Gen3G to calculate genes whose expression on the maternal side of the placenta was associated with offspring neurodevelopment (SDQ score at ages 3 and 5, ASEBA score at age 5). We used normalized (using base R scale) continuous neurodevelopment scores as the predictor, gene expression as the outcome, and included the following covariates: maternal age (normalized with base R scale), maternal education (categorical), delivery mode, fetal sex, and sequencing batch. We used consistent exclusion criteria as detailed above (“Hypoxia gene score correlations in Gen3G”). DESeq2 was used to calculate the association strength and significance. We calculated the associations based on on each ND testing done at 3y and/or 5y (SDQ 3, SDQ 5, and ASEBA 5),

### NDD score generation and application to sn/sc RNA seq

We used decoupleR (ulm) to calculate neurodevelopment impairment gene scores (NDIGSs) using the weights derived from the association described above, using the top 500 genes by p-value of association for each outcome. The test statistic from DESeq2 was used as the weights in decoupleR. We then applied these NDIGSs to the single nucleus (generated in this paper) and single cell (Cao et al.^28^) normalized expression matrices. This resulted in a gene score for each neurodevelopmental measure (SDQ 3, SDQ 5, and ASEBA 5) for each cell/ nucleus. We then grouped expression by cell type on the maternal side of the placenta in the single nucleus dataset, and all together in the Cao et al. dataset because side of origin was not available.

Next, we calculated a hypoxia gene score (HGS) similarly using decoupleR, with PROGENy as the gene sets. We performed Spearman correlations between the HGS and each NDIGS in each cell type.

We used the mediation 4.5.0^45^ package to calculate the average casual mediated effect (ACME) and average direct effect (ADE). The ACME is the estimated the indirect effect of BMI on NDIGS through HGS, and the ADE is the association between BMI and NDIGS that remains when accounting for HGS. BMI was the treatment variable and HGS was the mediator.

## Experimental Models and Subject Details

For demographics of the single-nucleus RNA seq cohort, please see Table S2. For demographics of the samples used from Gen3G in this study, please see Table S4.

## Data and code availability

The single-nucleus RNA-seq data generated in this study, and code used to analyze it, will be made publicly available upon publication. Single-cell RNA-seq data from Cao et al. is publicly available: GEO accession code: GSE156793. Gen3G is protected data available via application on dbGaP, accession code: phs003151.v1.p1.

## Author contributions

F.G.-R. and E.A.L. designed the study, with clinical guidance from R.M.R. and M.-F.H. F.G.-R. generated the single-nucleus RNA-seq data. F.G.-R. performed the analyses, with help from S.M. and F.W. F.G.-R. and S.M. had unrestricted access to all data. M.-F.H. supervised analyses with Gen3G data, and P.-E.J. supervised RNA-seq analyses from Gen3G data. R.M.R. and E.A.L. supervised single-nucleus RNA-seq analyses. F.G.-R. wrote the manuscript, with major edits from M.-F.H. and R.M.R. All authors agreed to submit the manuscript, read and approved the final draft and take full responsibility of its content, including the accuracy of the data.

## Acknowledgements

We thank the Women and Infants Health Specimen Consortium (WIHSC) Biobank at Washington University School of Medicine for the placental samples used in this study. We thank Diane Shao for her guidance on the frozen tissue handling and snRNA-seq data generation. This work was supported by the National Institute of Health (NIH) (DP2 AG072437), the Suh Kyungbae Foundation, and the Allen Discovery Center program, a Paul G. Allen Frontiers Group advised program of the Paul G. Allen Family Foundation. RNA sequencing in Gen3G was supported by a grant from the NIH (R01HD094150). Gen3G was initially supported by a Fonds de recherche du Québec – Santé (FRQS) operating grant (grant #20697); Canadian Institute of Health Research (CIHR) operating grants (grant #MOP 115071 and #PJT-152989). We thank all participants who provided placental samples referenced in this study.

