## Supplemental Figures and Tables for "Maternal obesity and offspring neurodevelopment are associated with hypoxic gene expression in term human placenta"

Figure S1: Proportions of each batch for each cell type

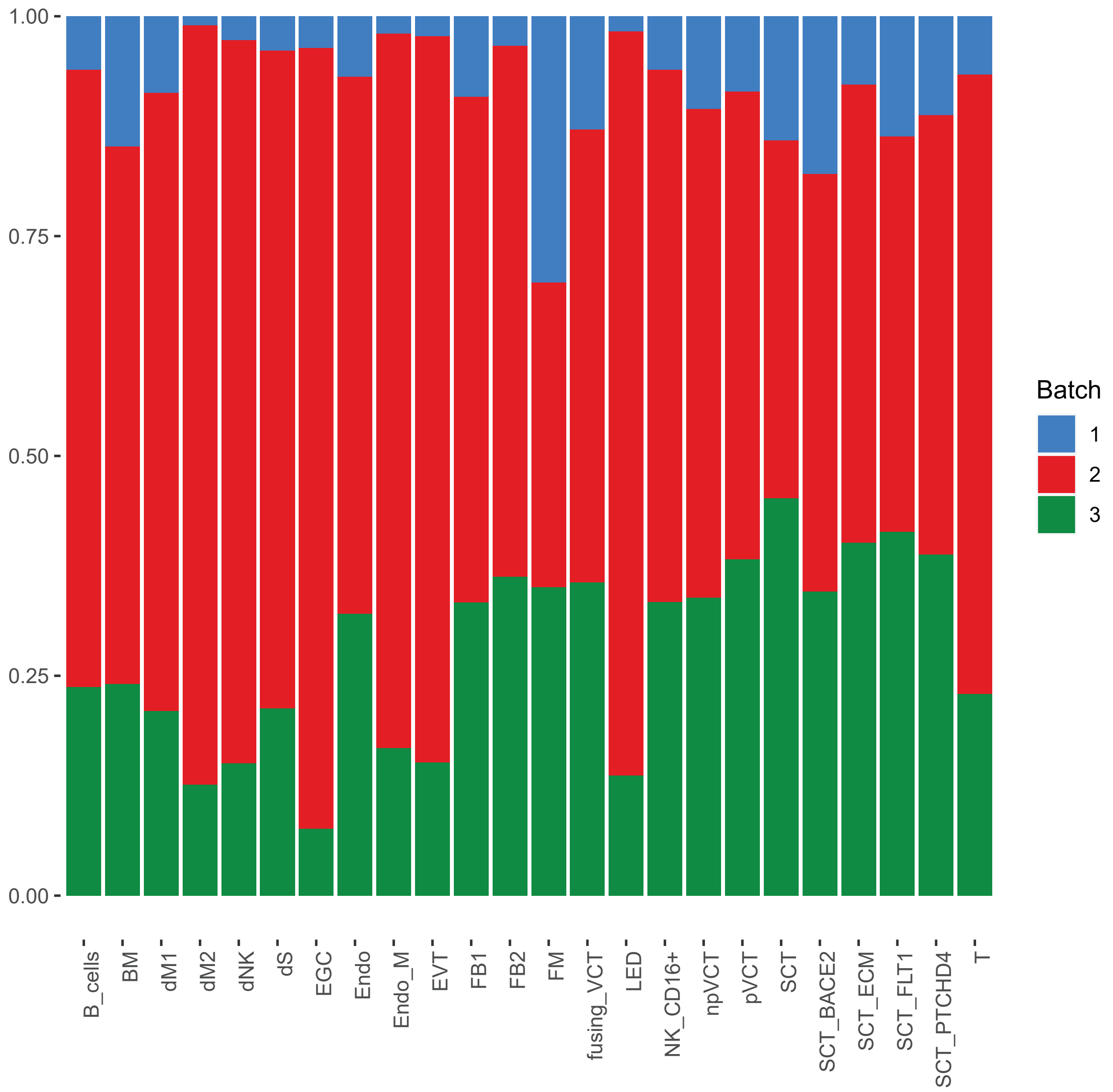

See main Figure 1 legend for abbreviations.

Figure S2: Proportions of each side (maternal facing or fetal facing) for each cell type. Side of origin is separate from person of origin (i.e. fetal or maternal DNA, as shown by cell type in Figure S3). Both sides have cell types of maternal and fetal origin.

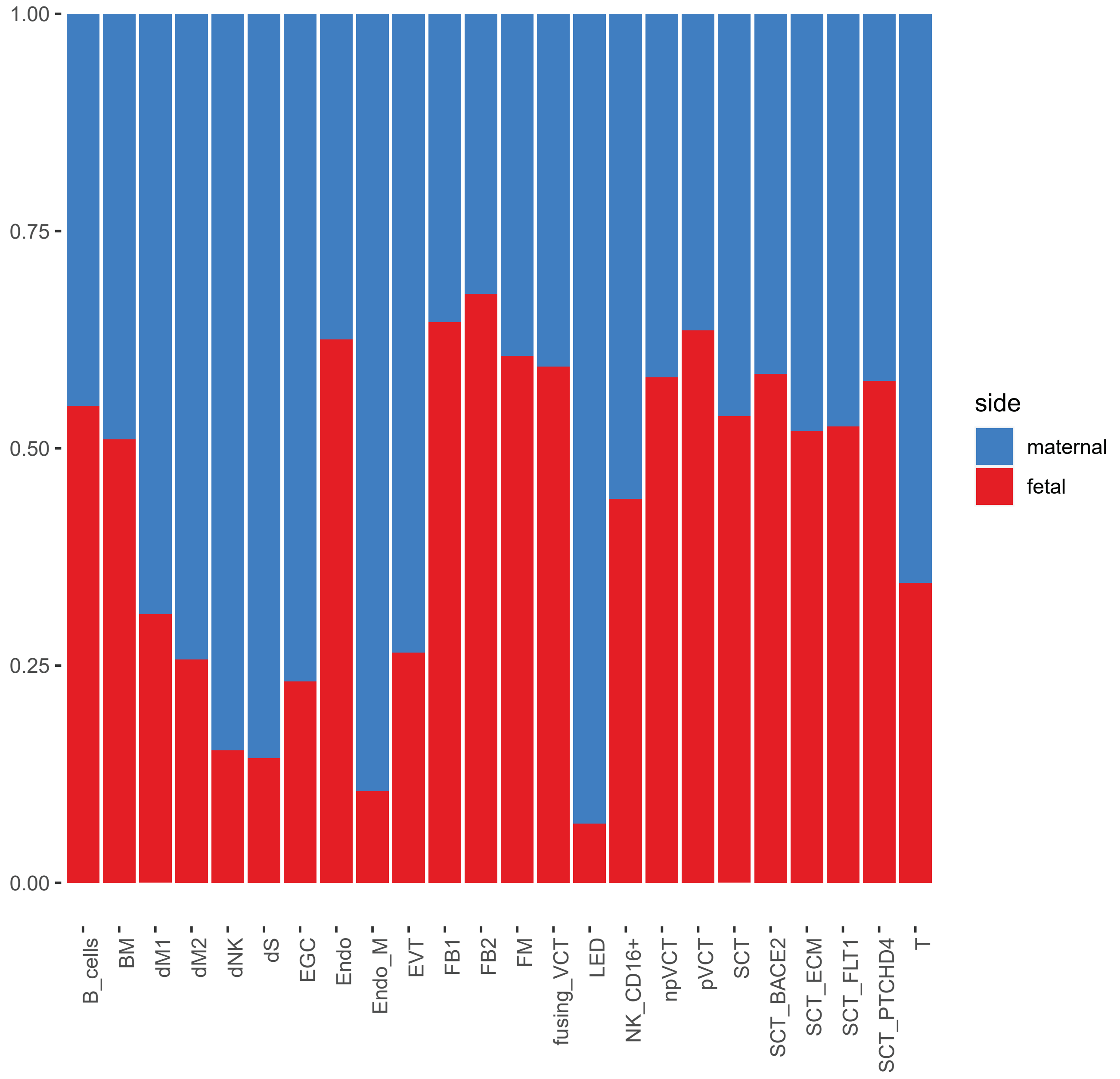

See main Figure 1 legend for abbreviations.

Figure S3: Inferred maternal/ fetal origin on cells based on sex-chromosome gene expression among male fetus placenta

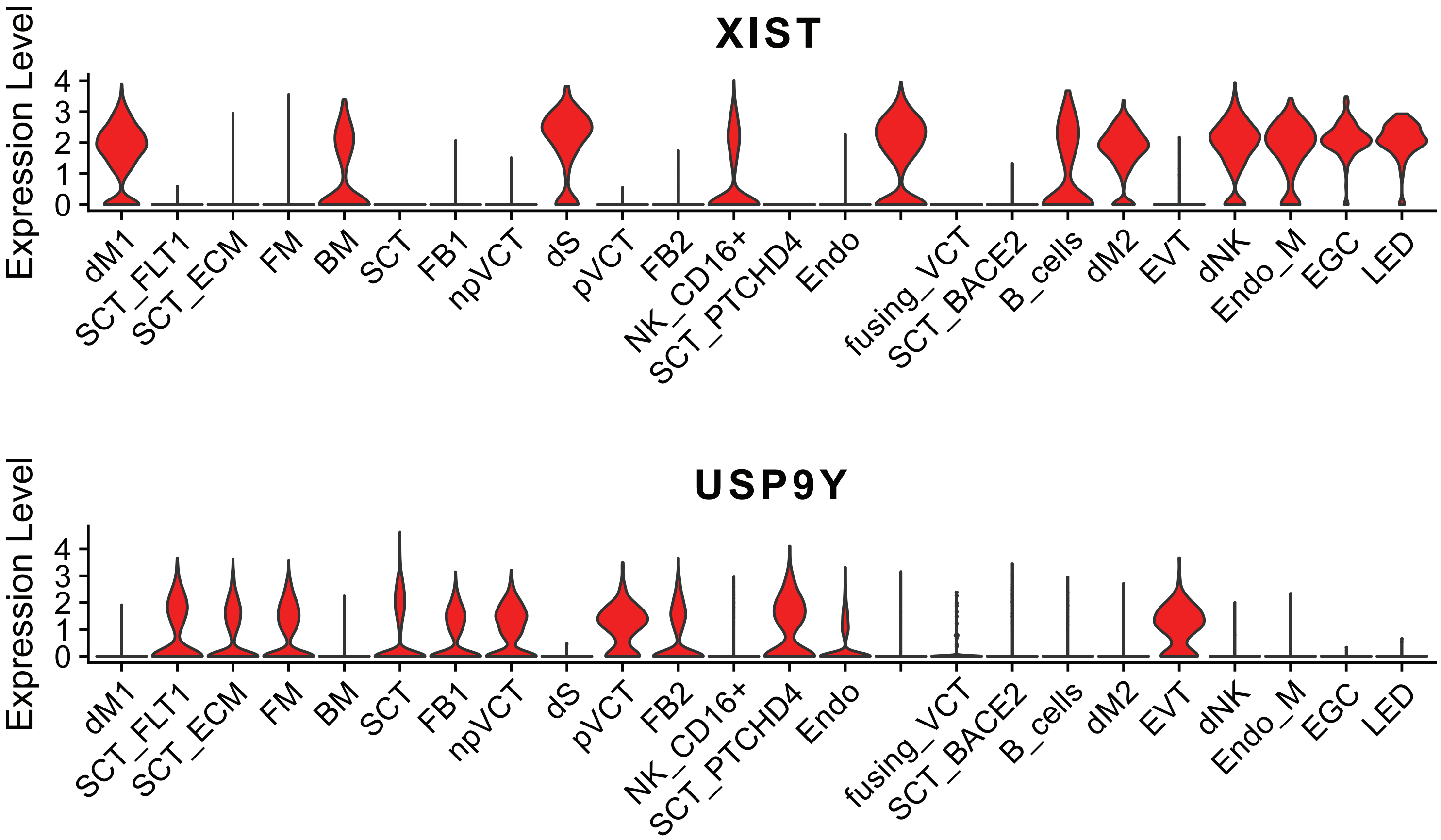

See main Figure 1 legend for abbreviations.

Figure S4: Gene ontology enrichment for markers of the SCT_ECM cell type. Driver terms from gProfiler are highlighted in blue.

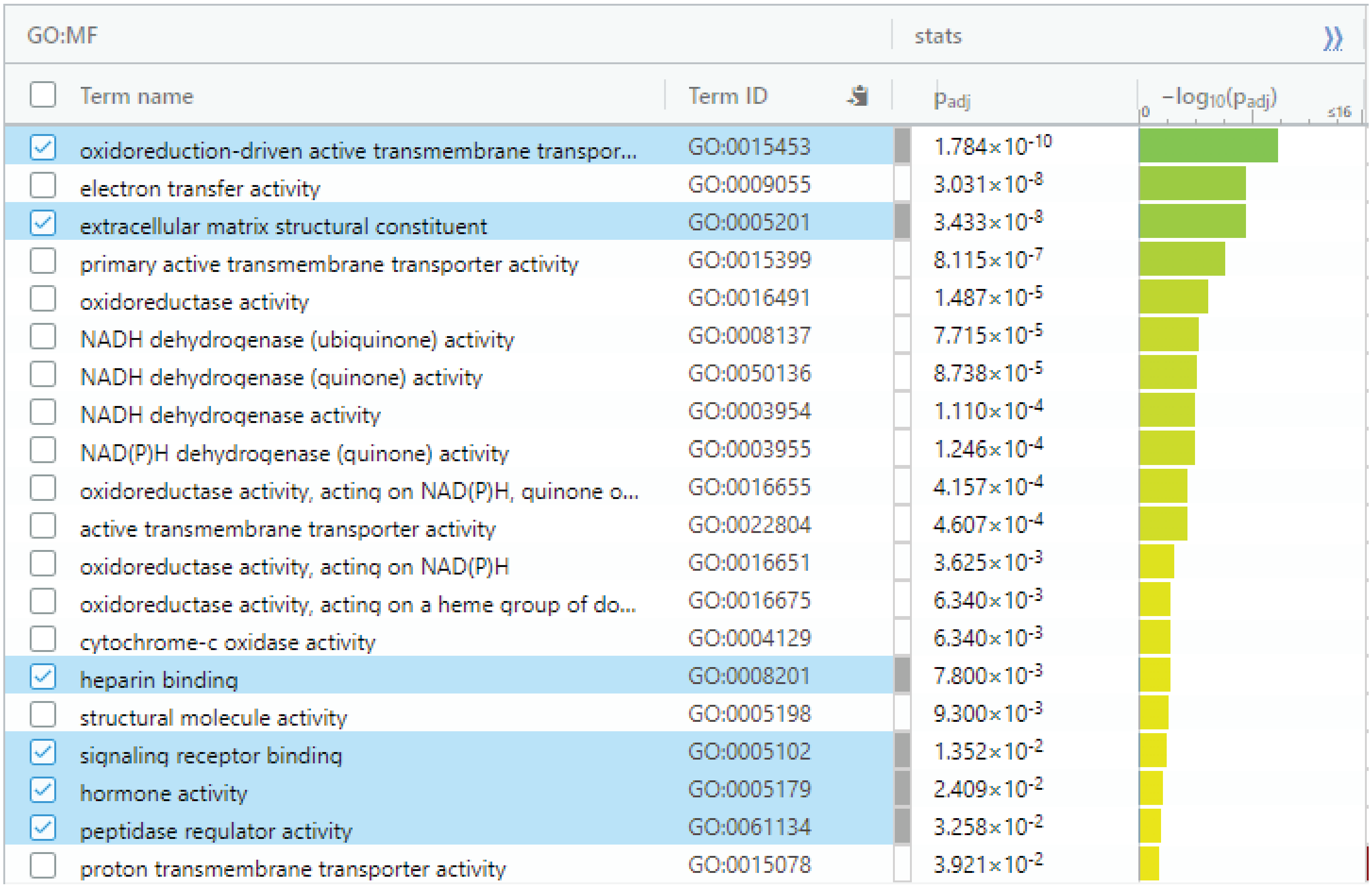

| GO:MF | | stats | |
|---|---|---|---|
| Term name | Term ID | $p_{adj}$ | $-\log_{10}(p_{adj})$ |
| oxidoreduction-driven active transmembrane transpor... | GO:0015453 | $1.784\times10^{-10}$ | |
| electron transfer activity | GO:0009055 | $3.031\times10^{-8}$ | |
| extracellular matrix structural constituent | GO:0005201 | $3.433\times10^{-8}$ | |
| primary active transmembrane transporter activity | GO:0015399 | $8.115\times10^{-7}$ | |
| oxidoreductase activity | GO:0016491 | $1.487\times10^{-5}$ | |
| NADH dehydrogenase (ubiquinone) activity | GO:0008137 | $7.715\times10^{-5}$ | |
| NADH dehydrogenase (quinone) activity | GO:0050136 | $8.738\times10^{-5}$ | |
| NADH dehydrogenase activity | GO:0003954 | $1.110\times10^{-4}$ | |
| NAD(P)H dehydrogenase (quinone) activity | GO:0003955 | $1.246\times10^{-4}$ | |
| oxidoreductase activity, acting on NAD(P)H, quinone o... | GO:0016655 | $4.157\times10^{-4}$ | |
| active transmembrane transporter activity | GO:0022804 | $4.607\times10^{-4}$ | |
| oxidoreductase activity, acting on NAD(P)H | GO:0016651 | $3.625\times10^{-3}$ | |
| oxidoreductase activity, acting on a heme group of do... | GO:0016675 | $6.340\times10^{-3}$ | |
| cytochrome-c oxidase activity | GO:0004129 | $6.340\times10^{-3}$ | |
| heparin binding | GO:0008201 | $7.800\times10^{-3}$ | |
| structural molecule activity | GO:0005198 | $9.300\times10^{-3}$ | |
| signaling receptor binding | GO:0005102 | $1.352\times10^{-2}$ | |
| hormone activity | GO:0005179 | $2.409\times10^{-2}$ | |
| peptidase regulator activity | GO:0061134 | $3.258\times10^{-2}$ | |
| proton transmembrane transporter activity | GO:0015078 | $3.921\times10^{-2}$ | |

Figure S5: Cell Type Proportions on the materal facing side by BMI

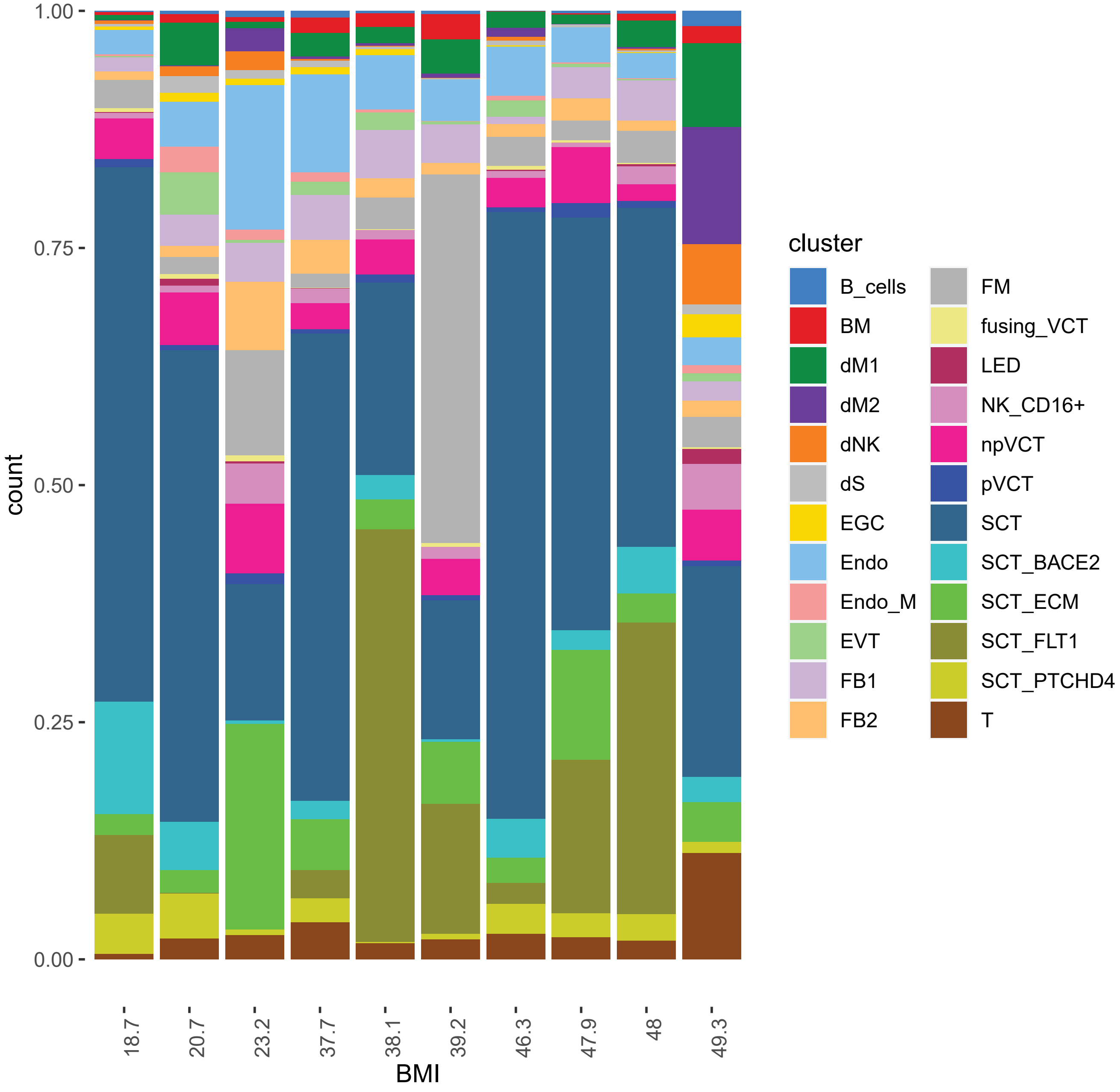

See main Figure 1 legend for abbreviations.

Fig S6: Cell Type Proportions on the fetal facing side by BMI

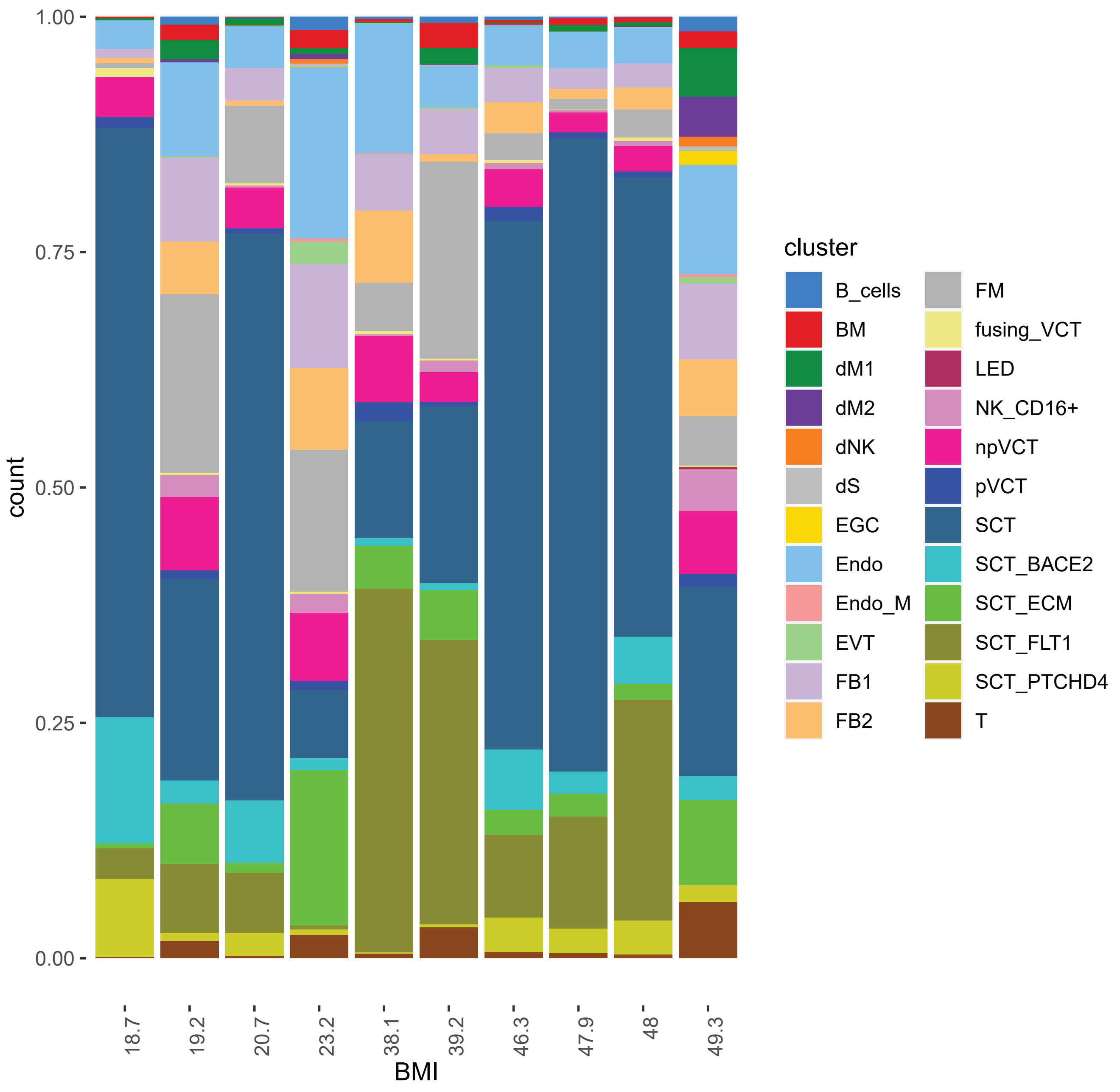

See main Figure 1 legend for abbreviations.

Figure S7: SDQ 5 and ASEBA 5 NDIGS for each nucleus in the term sn-RNA-seq dataset, grouped by cell type

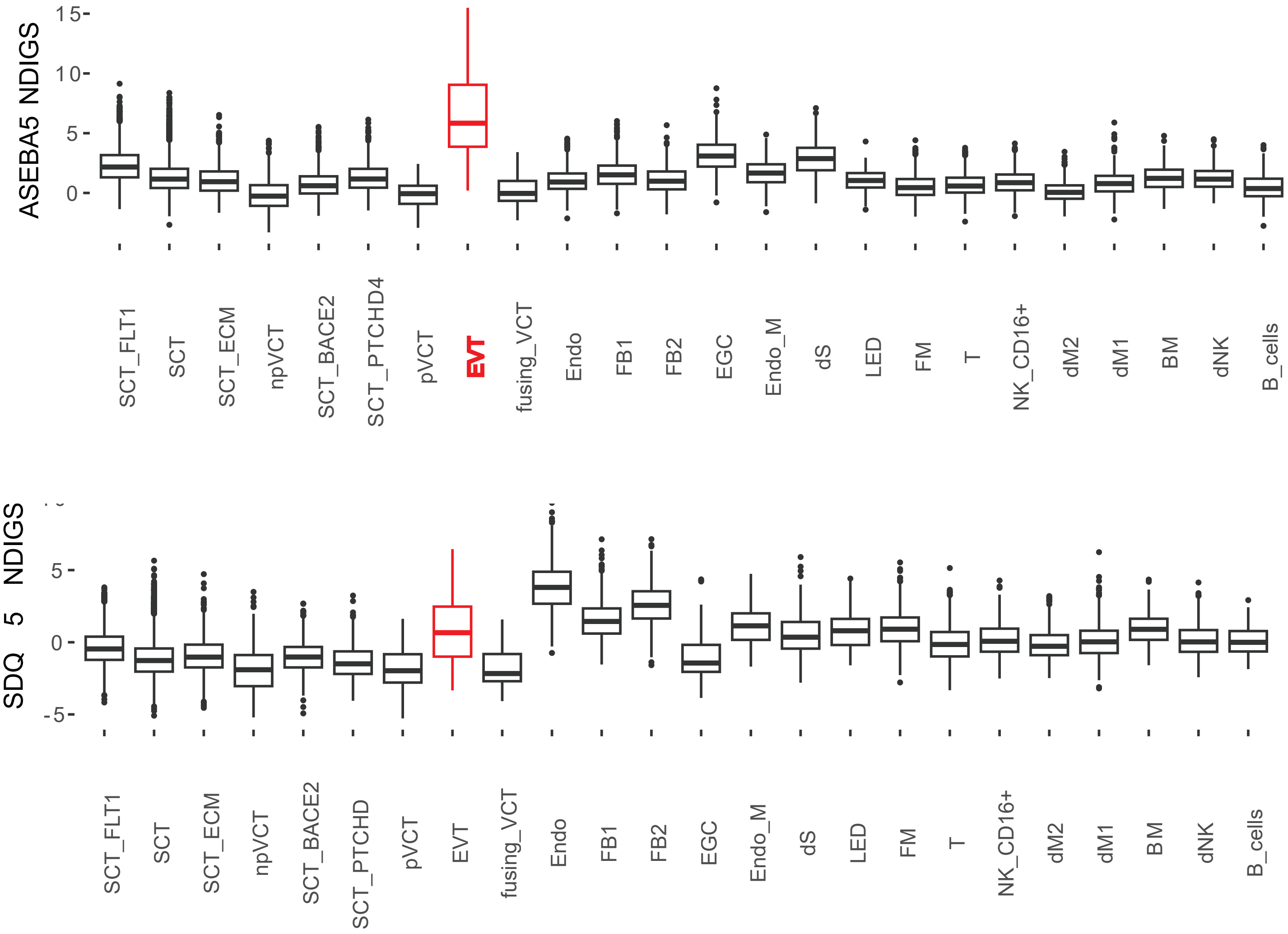

See main Figure 1 legend for abbreviations.

Figure S8: NDIGSs by cell type and BMI category (lean or maternal obesity)

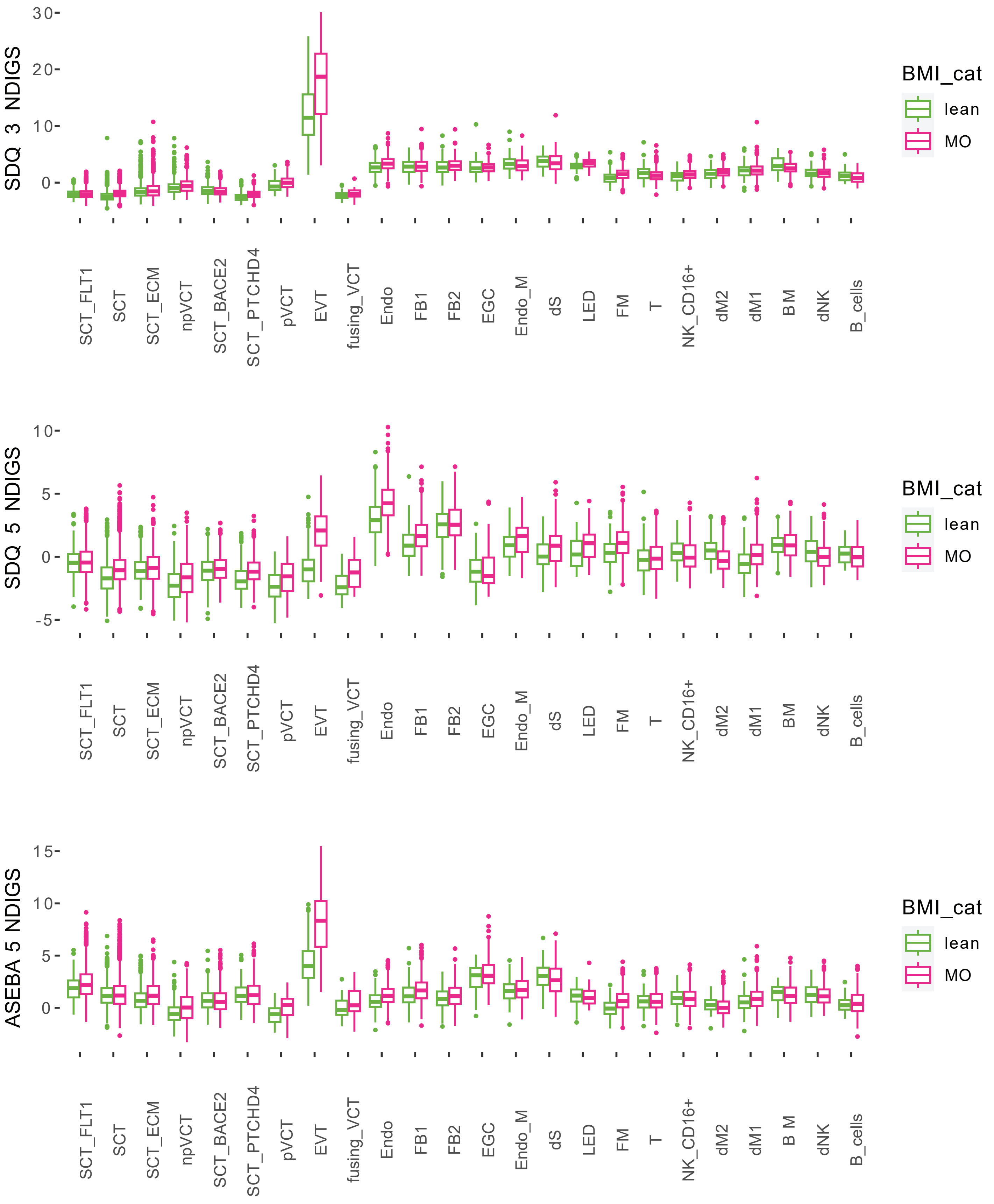

See main Figure 1 legend for abbreviations.

Figure S9: SDQ 5 and ASEBA 5 NDIGS for each cell the second trimester sc-RNA-seq dataset, grouped by cell type

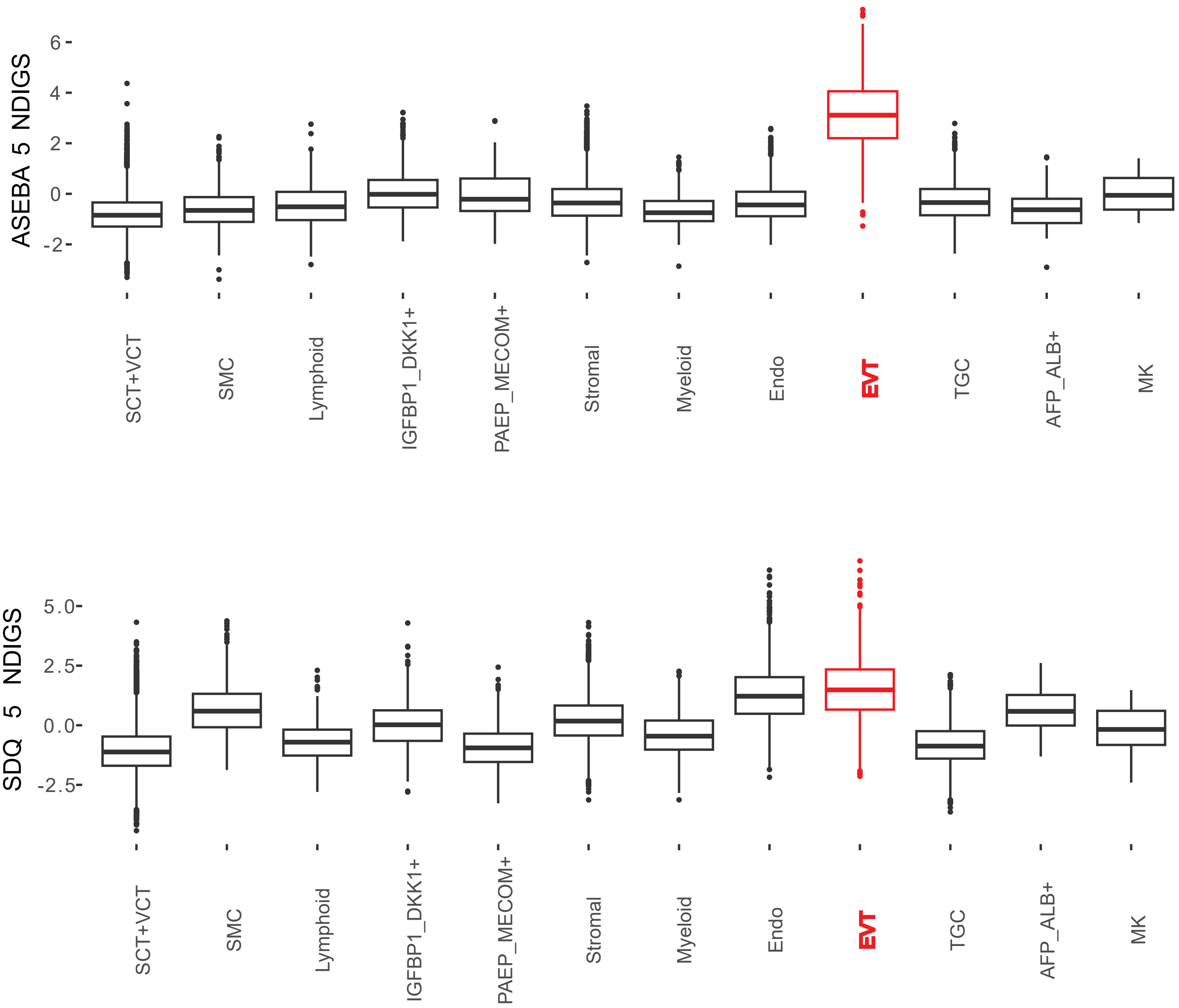

See main Figure 5 legend for abbreviations.

Table S1. Demographic charcteristics of placenta used in the single-nucleus cohort of data generated here.
VI= Vaginal Induction VS: Vaginal Spontaenous CS: Cesarean section

| Age at enrollment | Race | Infant gender | BMI | Delivery Mode | Included on maternal side | Included on fetal side |
|---|---|---|---|---|---|---|
| 30 | White | Female | 18.7 | VI | yes | yes |
| 25 | Black | Female | 20.7 | VS | yes | yes |
| 27 | Black | Male | 23.2 | CS | yes | yes |
| 19 | Black | Male | 19.2 | CS | no | yes |
| 29 | White | Female | 38.1 | VI | yes | yes |
| 22 | Black | Female | 39.2 | VS | yes | yes |
| 26 | Black | Female | 46.3 | VS | yes | yes |
| 20 | Black | Male | 37.7 | VS | yes | no |
| 30 | Black | Male | 49.3 | CS | yes | yes |
| 33 | White | Female | 48.0 | VI | yes | yes |
| 26 | Black | Female | 47.9 | VS | yes | yes |

Table S2. Enrichment of hypoxia associated genes among DEGs by maternal BMI in each cell type, and each placental face.

| Pathway | score | p_value | FDR | cell_type |
|---|---|---|---|---|
| Hypoxia | 7.04 | 2.09E-12 | 2.92E-11 | ms EVT |
| Hypoxia | 6.95 | 3.88E-12 | 5.44E-11 | fs SCT_FLT1 |
| Hypoxia | 6.45 | 1.13E-10 | 1.58E-09 | fs SCT |
| Hypoxia | 5.14 | 2.71E-07 | 3.80E-06 | fs SCT_ECM |
| Hypoxia | 3.75 | 1.79E-04 | 2.51E-03 | fs npVCT |
| Hypoxia | 3.41 | 6.50E-04 | 9.09E-03 | ms FM |
| Hypoxia | 3.29 | 1.01E-03 | 1.42E-02 | ms dM2 |
| Hypoxia | 3.27 | 1.09E-03 | 7.62E-03 | ms dM1 |
| Hypoxia | 3.23 | 1.22E-03 | 8.54E-03 | ms SCT |
| Hypoxia | 3.17 | 1.55E-03 | 0.0109 | fs FM |
| Hypoxia | 2.97 | 2.98E-03 | 0.0139 | ms Endo |
| Hypoxia | -2.90 | 3.77E-03 | 0.0264 | ms dNK |
| Hypoxia | -2.87 | 4.12E-03 | 0.0289 | ms Endo_M |
| Hypoxia | -2.75 | 5.94E-03 | 0.0416 | ms BM |
| Hypoxia | 2.75 | 5.94E-03 | 0.0832 | fs NK_CD16+ |
| Hypoxia | 2.67 | 7.65E-03 | 0.0860 | fs SCT_PTCHD4 |
| Hypoxia | 2.58 | 0.0100 | 0.0279 | fs Endo |
| Hypoxia | 2.53 | 0.0116 | 0.0324 | fs pVCT |
| Hypoxia | 2.49 | 0.0127 | 0.0295 | fs FB1 |
| Hypoxia | 2.48 | 0.0131 | 0.1833 | ms SCT_FLT1 |
| Hypoxia | -2.48 | 0.0132 | 0.0307 | ms SCT_BACE2 |
| Hypoxia | 2.08 | 0.0372 | 0.1302 | fs dM1 |
| Hypoxia | 2.05 | 0.0406 | 0.1896 | ms SCT_ECM |
| Hypoxia | 2.03 | 0.0421 | 0.0983 | ms FB1 |
| Hypoxia | -1.66 | 0.0968 | 0.2710 | ms dS |
| Hypoxia | 1.40 | 0.1605 | 0.4374 | fs T |
| Hypoxia | -1.21 | 0.2258 | 0.5268 | ms FB2 |
| Hypoxia | 1.09 | 0.2761 | 0.5522 | ms npVCT |
| Hypoxia | 0.93 | 0.3538 | 0.4503 | fs FB2 |
| Hypoxia | 0.81 | 0.4192 | 0.5868 | ms pVCT |
| Hypoxia | -0.43 | 0.6674 | 0.7188 | ms T |
| Hypoxia | 0.41 | 0.6786 | 0.9188 | fs BM |
| Hypoxia | -0.37 | 0.7083 | 0.8936 | ms SCT_PTCHD4 |
| Hypoxia | 0.32 | 0.7456 | 0.8611 | fs fusing_VCT |
| Hypoxia | 0.25 | 0.8060 | 0.8932 | ms B_cells |
| Hypoxia | -0.20 | 0.8384 | 0.8384 | ms NK_CD16+ |
| Hypoxia | -0.08 | 0.9370 | 0.9667 | fs SCT_BACE2 |
| Hypoxia | -0.03 | 0.9765 | 0.9765 | fs B_cells |
| Hypoxia | 0.03 | 0.9786 | 0.9786 | ms fusing_VCT |

Table S3 Test statistics of hypoxia associated genes in the four cell types whose DEGs by BMI had the strongest enrichment for hypoxia response genes.

| | ms EVT | fs SCT | fs SCT_FLT1 | fs SCT_ECM |
|---|---|---|---|---|
| AFAP1 | 4.58 | NA | NA | NA |
| MTRNR2L12 | 4.21 | NA | NA | NA |
| GSE1 | 3.91 | NA | NA | NA |
| MALAT1 | -3.68 | NA | NA | NA |
| DIP2C | 3.45 | NA | NA | NA |
| AC106729.1 | 3.39 | NA | NA | NA |
| PPARG | 3.28 | NA | NA | NA |
| MME | 3.22 | NA | NA | NA |
| CRISPLD2 | 3.15 | NA | NA | NA |
| MICAL3 | 3.09 | NA | NA | NA |
| KDM4B | 1.37 | 3.90 | 2.32 | 2.29 |
| HK2 | 2.21 | 3.15 | 1.76 | 0.00 |
| PNRC1 | -0.23 | 3.07 | 3.30 | 1.15 |
| GBE1 | 0.44 | -3.01 | -0.64 | 0.83 |
| SEMA4B | 0.88 | 2.99 | 2.52 | 2.40 |
| PLIN2 | 1.62 | 2.90 | 1.78 | 2.36 |
| ARID3A | 2.26 | 2.76 | 2.37 | 0.97 |
| NFIL3 | 0.30 | 2.72 | 2.18 | 1.86 |
| PFKFB4 | 1.96 | 2.69 | 2.27 | 1.56 |
| PDK3 | -0.35 | 2.55 | 1.42 | 0.78 |
| SERTAD2 | -0.18 | 1.27 | 3.50 | 0.39 |
| PPME1 | -1.17 | -1.09 | -3.32 | 0.02 |
| ERO1A | 0.07 | 1.07 | 2.99 | -0.25 |
| FOSL2 | 0.80 | 2.03 | 2.54 | 1.39 |
| COASY | 0.19 | -0.34 | -2.51 | 1.25 |
| ENO1 | 0.38 | 0.79 | 2.49 | -0.08 |
| ADM | 0.05 | 2.36 | 2.47 | 1.96 |
| GPI | 0.59 | 0.77 | 2.45 | 0.49 |
| PPP1R13L | -0.72 | 1.15 | 0.94 | 2.36 |
| SLC6A6 | 0.62 | 1.89 | 1.43 | 2.12 |
| SLC35F6 | 0.14 | -0.05 | 1.34 | -1.95 |
| LARP4 | 0.39 | 2.20 | 2.22 | 1.91 |
| IPMK | -1.19 | 1.17 | 1.00 | 1.75 |

NAs come from genes that were not included in the differential expression testing because their expression was too low (<10 counts total across all samples).

Table S4. Demographic charcteristics of placental samples from Gen3G used in this study

| | samples form maternal facing side | samples from fetal facing side |
|---|---|---|
| N | 262 | 83 |
| Of European descent | 98% | 99% |
| Maternal Age | 28.7 [25.6-31.4] | 28.7 [25.1-31.2] |
| Fetal Sex | 47% female | 46% female |
| Delivery Mode | 96% vaginal | 95% vaginal |
| Apgar 5 minutes | 9 [9-10] | 9 [9-10] |
| Cord Blood pH | 7.26 [7.20-7.30] | 7.26 [7.20-7.31] |
| Hemoglobin | 155 [145-168] | 154 [145-164] |
| Birthweight | 3465 [3225-3725] | 3470 [3225-3705] |

Values represent median [interquartile range] or percentage.

Table S5. Results of Spearman correlation between hypoxia gene score and offspring measures.

| category | test | side | cor | p |
|---|---|---|---|---|
| Acute | APGAR at 5 min | fs | -0.34373 | 3.09E-04 |
| Acute | APGAR at 5 min | ms | -0.12334 | 4.70E-02 |
| Acute | Cord Blood pH | fs | -0.41313 | 2.87E-05 |
| Acute | Cord Blood pH | ms | -0.24713 | 8.10E-05 |
| Acute | Hemoglobin | fs | 0.317709 | 8.54E-04 |
| Acute | Hemoglobin | ms | 0.139822 | 2.42E-02 |
| Growth | Birthweight | fs | -0.19659 | 4.24E-02 |
| Growth | Birthweight | ms | 0.008753 | 8.88E-01 |
| Growth | BMI at age 3 | fs | 0.045488 | 7.10E-01 |
| Growth | BMI at age 3 | ms | 0.185451 | 2.57E-02 |
| Growth | BMI at age 5 | fs | 0.039431 | 7.25E-01 |
| Growth | BMI at age 5 | ms | 0.028842 | 7.01E-01 |
| NDD | ASEBA at age 5 | fs | 0.027531 | 8.10E-01 |
| NDD | ASEBA at age 5 | ms | 0.167249 | 2.74E-02 |
| NDD | SDQ at age 3 | fs | 0.17402 | 1.53E-01 |
| NDD | SDQ at age 3 | ms | 0.25615 | 1.50E-03 |
| NDD | SDQ at age 5 | fs | 0.009676 | 9.30E-01 |
| NDD | SDQ at age 5 | ms | 0.181345 | 1.38E-02 |

Table S6: Correlations between neurodevelopmental gene scores (NDIGSs) and hypoxia gene score (HGS) for each cell type in the term placenta sn-RNA-seq dataset

| Cell Type | SDQ 3 cor | SDQ 3 p | SDQ 5 cor | SDQ 5 p | ASEBA 5 cor | ASEBA 5 p |
|---|---|---|---|---|---|---|
| EVT | 0.537 | <2.2e-16 | 0.550 | <2.2e-16 | 0.462 | <2.2e-16 |
| Endo_M | 0.392 | 1.06E-09 | 0.357 | 3.52E-08 | 0.067 | 3.10E-01 |
| dS | 0.388 | 2.28E-08 | 0.417 | 1.51E-09 | -0.021 | 7.69E-01 |
| Endo | 0.314 | 3.22E-39 | 0.003 | 8.94E-01 | 0.048 | 5.11E-02 |
| EGC | 0.251 | 2.26E-04 | 0.212 | 1.97E-03 | 0.074 | 2.84E-01 |
| FM | 0.221 | 9.04E-19 | 0.124 | 7.37E-07 | 0.006 | 8.22E-01 |
| FB1 | 0.214 | 1.60E-11 | -0.002 | 9.57E-01 | -0.105 | 9.68E-04 |
| dNK | 0.194 | 1.50E-04 | 0.062 | 2.31E-01 | -0.092 | 7.32E-02 |
| dM2 | 0.172 | 1.61E-05 | -0.052 | 1.97E-01 | -0.106 | 7.94E-03 |
| FB2 | 0.137 | 4.77E-04 | -0.033 | 4.01E-01 | -0.039 | 3.25E-01 |
| BM | 0.128 | 2.46E-02 | 0.031 | 5.93E-01 | -0.027 | 6.35E-01 |
| SCT_BACE2 | 0.110 | 3.03E-04 | 0.084 | 6.13E-03 | 0.072 | 1.82E-02 |
| pVCT | 0.099 | 1.48E-01 | 0.162 | 1.67E-02 | -0.026 | 7.05E-01 |
| dM1 | 0.094 | 4.39E-03 | 0.124 | 1.72E-04 | -0.034 | 3.01E-01 |
| LED | 0.088 | 3.60E-01 | 0.188 | 5.03E-02 | 0.023 | 8.14E-01 |
| T | 0.076 | 1.48E-02 | -0.059 | 5.91E-02 | -0.208 | 2.35E-11 |
| NK_CD16+ | 0.070 | 1.03E-01 | 0.001 | 9.84E-01 | -0.206 | 1.51E-06 |
| npVCT | 0.028 | 3.17E-01 | 0.026 | 3.63E-01 | -0.120 | 2.35E-05 |
| B_cells | -0.015 | 8.51E-01 | 0.257 | 1.23E-03 | -0.015 | 8.52E-01 |
| fusing_VCT | -0.032 | 7.76E-01 | -0.131 | 2.39E-01 | -0.205 | 6.54E-02 |
| SCT_ECM | -0.032 | 1.94E-01 | 0.215 | 1.54E-18 | 0.008 | 7.51E-01 |
| SCT_FLT1 | -0.117 | 1.84E-12 | 0.277 | 7.72E-64 | 0.143 | 7.09E-18 |
| SCT | -0.129 | 2.28E-41 | 0.180 | 6.41E-79 | -0.018 | 6.29E-02 |
| SCT_PTCHD4 | -0.213 | 3.50E-08 | 0.177 | 5.30E-06 | -0.062 | 1.12E-01 |

Table S7: Correlations between NDIGSs and HGS for each cell type in the second trimester placenta sc-RNA-seq dataset

| Cell Type | SDQ 3 cor | SDQ 3 p | SDQ 5 cor | SDQ 5 p | ASEBA 5 cor | ASEBA 5 p |
|---|---|---|---|---|---|---|
| EVT | 0.285 | 3.60E-30 | 0.243 | 4.07E-22 | 0.175 | 5.72E-12 |
| MK | 0.285 | 1.27E-01 | 0.152 | 4.20E-01 | -0.067 | 7.24E-01 |
| IGFBP1_DKK1+ | 0.193 | 2.61E-07 | 0.076 | 4.49E-02 | 0.069 | 6.87E-02 |
| Stromal | 0.190 | 9.77E-69 | 0.079 | 5.04E-13 | -0.020 | 6.69E-02 |
| PAEP_MECOM+ | 0.168 | 4.30E-03 | 0.011 | 8.54E-01 | 0.124 | 3.58E-02 |
| SCT+VCT | 0.154 | 3.84E-78 | 0.172 | 2.24E-97 | 0.067 | 4.66E-16 |
| Endo | 0.145 | 2.65E-08 | 0.061 | 1.98E-02 | 0.015 | 5.79E-01 |
| SMC | 0.127 | 1.66E-04 | 0.081 | 1.63E-02 | -0.048 | 1.57E-01 |
| Lymphoid | 0.104 | 2.53E-02 | 0.076 | 1.01E-01 | 0.037 | 4.28E-01 |
| Myeloid | 0.043 | 4.30E-01 | 0.043 | 4.26E-01 | 0.065 | 2.33E-01 |
| TGC | 0.033 | 2.62E-01 | 0.115 | 1.01E-04 | 0.061 | 3.85E-02 |
| AFP_ALB+ | -0.028 | 7.98E-01 | 0.036 | 7.37E-01 | 0.013 | 9.03E-01 |
